# High-resolution spatial multi-omics reveals cell-type specific nuclear compartments

**DOI:** 10.1101/2023.05.07.539762

**Authors:** Yodai Takei, Yujing Yang, Jonathan White, Jina Yun, Meera Prasad, Lincoln J Ombelets, Simone Schindler, Long Cai

## Abstract

The mammalian nucleus is compartmentalized by diverse subnuclear structures. These subnuclear structures, marked by nuclear bodies and histone modifications, are often cell-type specific and affect gene regulation and 3D genome organization^1–3^. Understanding nuclear organization requires identifying the molecular constituents of subnuclear structures and mapping their associations with specific genomic loci in individual cells, within complex tissues. Here, we introduce two-layer DNA seqFISH+, which allows simultaneous mapping of 100,049 genomic loci, together with nascent transcriptome for 17,856 genes and a diverse set of immunofluorescently labeled subnuclear structures all in single cells in cell lines and adult mouse cerebellum. Using these multi-omics datasets, we showed that repressive chromatin compartments are more variable by cell type than active compartments. We also discovered a single exception to this rule: an RNA polymerase II (RNAPII)-enriched compartment was associated with long, cell-type specific genes (> 200kb), in a manner distinct from nuclear speckles. Further, our analysis revealed that cell-type specific facultative and constitutive heterochromatin compartments marked by H3K27me3 and H4K20me3 are enriched at specific genes and gene clusters, respectively, and shape radial chromosomal positioning and inter-chromosomal interactions in neurons and glial cells. Together, our results provide a single-cell high-resolution multi-omics view of subnuclear compartments, associated genomic loci, and their impacts on gene regulation, directly within complex tissues.

## Main

Recent imaging-based genome-wide multimodal technologies have enabled direct profiling of the 3D organization of the nucleus in single cells^4–9^, providing spatial context to our understanding of nuclear architecture derived from genome-wide sequencing-based approaches^10–15^. For example, we had previously shown that specific associations between genomic loci and subnuclear structures are conserved across single cells, despite the apparent variability in the 3D genome structure of individual cells^7, 8^. Furthermore, imaging-based transcriptomic approaches have revealed the organization of the nascent transcriptome within the nucleus^16^ and identified cell-type specific transcriptional programs in tissues^17^. However, due to optical crowding, current genome-wide imaging approaches are limited to resolving genomic sites at the megabase level^5–8^, limiting the achievable level of insight to larger genomic regions, rather than individual genes across the genome.

To enable a more detailed understanding of nuclear organization and its relationship to gene regulation, higher genomic resolution measurements are required together with diverse subnuclear structures in single cells. Here we introduce a two-layer barcoding DNA seqFISH+ scheme that increases the multiplexing capability of single-cell multi-omics to ∼100,000 species, up from the previous ∼10,000^16–18^, corresponding to 25-kb coverage across the genome (Methods) (Fig. 1b, Extended Data Fig. 3a). This approach enables the simultaneous investigation of gene expression profiles and chromatin organization at the sub-megabase resolution across the genome in individual cells (Fig. 1a, Extended Data Figs. 1-4), and enables identification of chromatin compartments and their associated genes in a cell-type specific fashion.

**Fig. 1.**
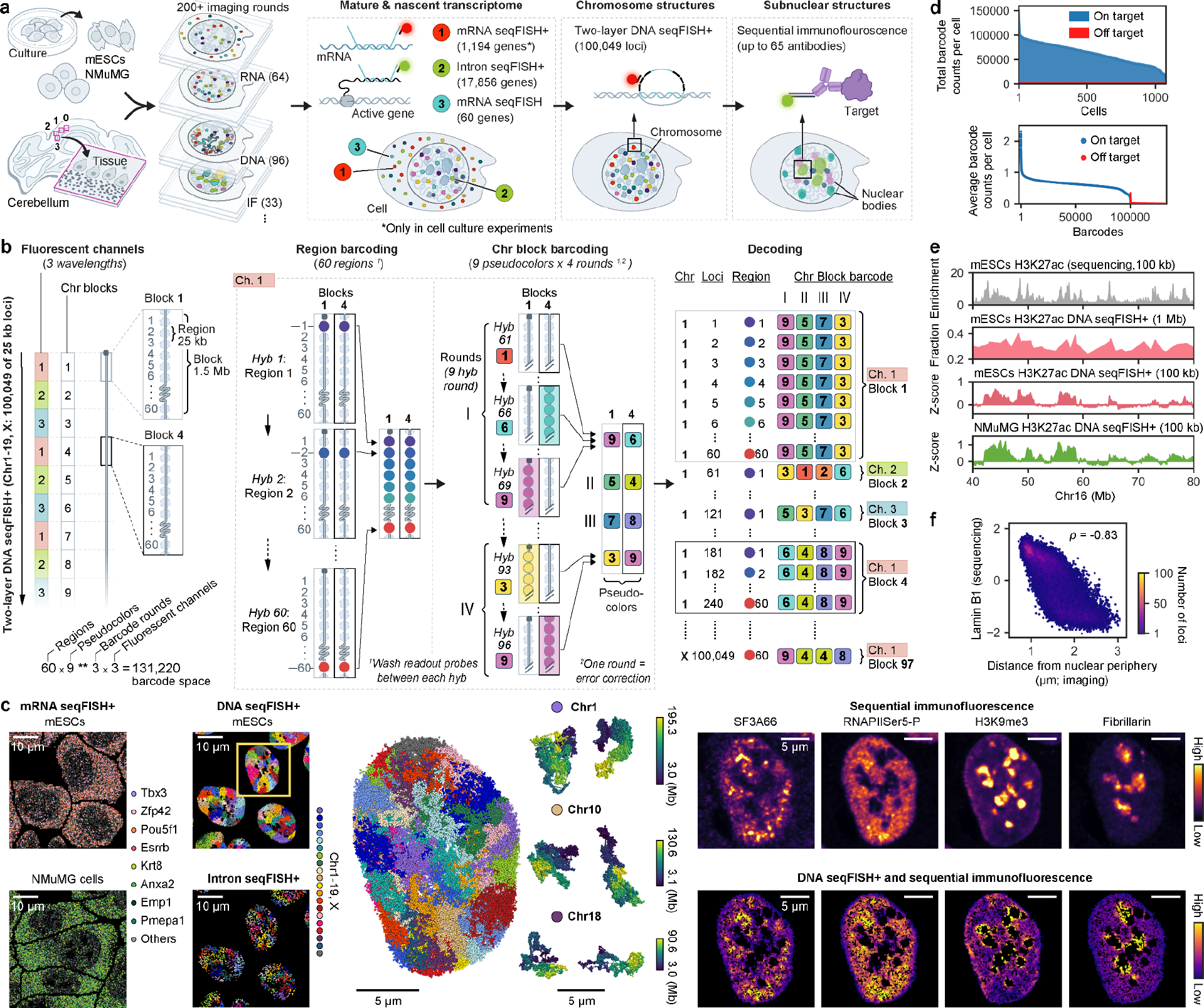
**Development and validation of single-cell spatial multi-omics technology. a**, Schematic of spatial multi-omics, imaging mature and nascent transcriptome as well as chromosome and subnuclear structures within the same single cells. **b**, Detailed schematic of DNA seqFISH+. Chromosomes are splitted into 1.5-Mb blocks, in which each 25-kb locus is imaged as a diffraction limited spot during the first 60 rounds of imaging with three orthogonal fluorescent channels. Then each 1.5-Mb block is uniquely barcoded across 4 rounds with 9-pseudocolors during the subsequent 36 rounds of imaging. Finally, by decoding the region and block identities, 100,049 of 25-kb loci can be uniquely resolved across 20 chromosomes. **c**, Visualization of decoded RNA (left) and DNA spots (middle), immunofluorescence raw images (top right), and DNA spots colored by z-score normalized immunofluorescence intensity (bottom right) in the nucleus from mESCs. **d**, Total on- and off-target barcode counts per cell (top). Average counts of individual barcode per cell (bottom). *n* = 100,049 and 31,171 DNA loci for on- and off-target barcodes. **e**, **f**, Comparison of our imaging-based chromatin profiles with those from previous DNA seqFISH+^7^ and sequencing-based technologies for the selected genomic region (**e**) and across the genome (**f**) with 100 kb binning (*n* = 25,110 loci). *n* = 1,076 cells from two biological replicates of mESCs and *n* = 384 cells from one biological replicate of NMuMG cells in **c**-**f**.

**Fig. 3.**
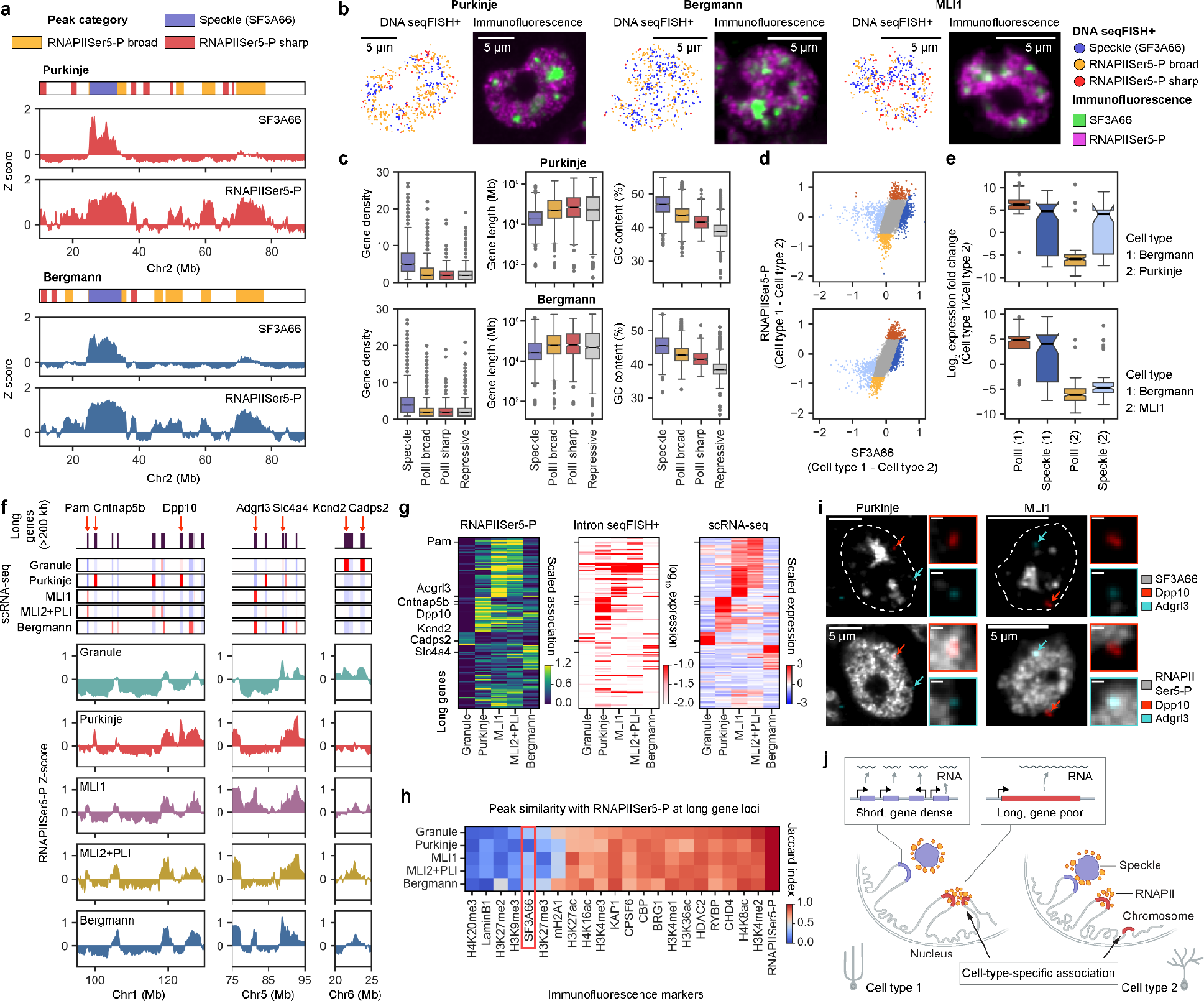
**Distinct subnuclear compartments organize transcriptionally-active genomic loci. a**, Representative classification of three types of active domains (nuclear speckle, RNAPIISer5-P broad and sharp) in Purkinje cells and Bergmann glia along with the chromatin profiling of SF3A66 (nuclear speckle marker) and RNAPIISer5-P. **b**, Visualization of genomic loci colored by the ensemble-average active domain classification in **a** (left) and corresponding raw immunofluorescence image (right) from a single z-section for each cell type. **c**, Comparison of genomic features across different active domains in each cell type. *n* = 1,405, 4,102, 679, 4,892 loci (Purkinje) and 3,161, 3,115, 411, 4,080 loci (Bergmann) from left to right category. **d**, Comparison of differential association of genomic loci with SF3A66 and RNAPIISer5-P between pairs of cell types. **e**, Comparison of differential mRNA expression between pairs of cell types at differentially associated loci for either PolII (RNAPIISer5-P) or Speckle (SF3A66) in each cell type, colored in **d**. *n* = 110, 55, 121, 94 loci (top) and 134, 94, 109, 67 loci (bottom) from left to right category. In box plots, the center lines for the median, boxes for the interquartile range, whiskers for values within 1.5 times the interquartile range, and points for outliers (**c**, **e**). **f**, Representative genomic regions with long genes (>200 kb) (top), corresponding mRNA expression^26^ (middle), and RNAPIISer5-P chromatin profiles (bottom) in each cell type. **g**, Heatmaps for RNAPIISer5-P chromatin profiles (left), nascent RNA expression (middle), and mRNA expression^26^ (right) for highly correlated long genes (*n* = 132 genes, Methods). **h**, Similarity of RNAPIISer5-P peaks with other markers on the long genes in **g** in each cell type. **i**, Representative single cell visualization of long gene loci with cell-type specific gene expression (Dpp10 in Purkinje cells and Adgrl3 in MLI1), relative to nuclear speckles (SF3A66) and RNAPIISer5-P with a maximum z-projection of two z-sections. Scale bars, 500 nm. **j**, Illustration showing nuclear speckle and RNAPIISer5-P subnuclear compartments associated with distinct genomic loci in a cell-type specific fashion. 200 kb binning (*n* = 12,562 loci in total) was used for the analysis and visualization (**a**, **c**-**h**). *n* = 2,336, 128, 263, 88, and 518 cells for Granule, Purkinje, MLI1, MLI2+PLI, and Bergmann glia cells from two biological replicates of the adult mouse cerebellum in **a**-**i**.

**Fig. 4.**
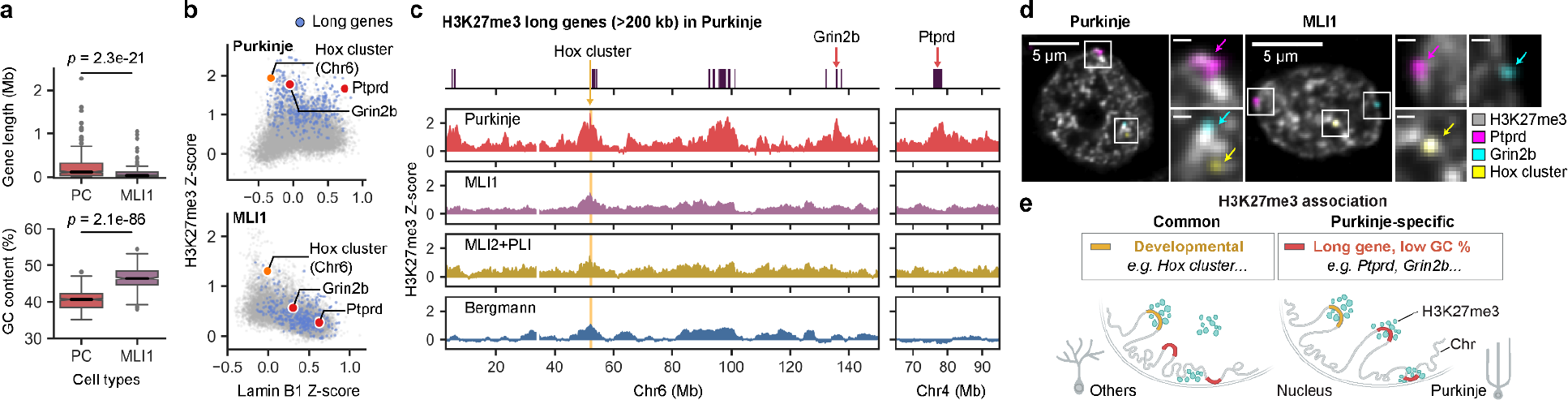
**Cell-type specific organization of H3K27me3 repressive subnuclear compartments. a**, Comparison of genomic features between loci that were differentially associated with H3K27me3 in two cell types. *n* = 600, 260 loci for Purkinje cells (PC) and MLI1. *p* values by two-sided Wilcoxon’s signed rank-sum test. The center lines for the median, boxes for the interquartile range, whiskers for values within 1.5 times the interquartile range, and points for outliers. **b**, Comparison of Lamin B1 and H3K27me3 chromatin profiles in each cell type. **c**, H3K27me3 profiles across cell types, highlighted with Hox cluster. Identified Purkinje-specific long gene loci associated with H3K27me3 are shown as blue dots (**b**) and binary heatmap (**c**). **d**, Representative single cell visualization of genomic loci highlighted (**b**, **c**), relative to H3K27me3. Scale bars, 500 nm. **e**, Illustration showing H3K27me3 subnuclear compartments associated with common and Purkinje-specific genomic loci at the nuclear interior or periphery. 200 kb binning (*n* = 12,562 loci in total) was used for the analysis and visualization (**a**-**c**). *n* = 128, 263, 88, and 518 cells for Purkinje, MLI1, MLI2+PLI, and Bergmann glia cells from two biological replicates of the adult mouse cerebellum in **a**-**d**.

### Two-layer seqFISH+: Single-cell spatial multi-omics technology

Building upon our previous multimodal technology^7, 8^, our goal was to employ a multi-omics approach to characterize the genomic landscape of individual cells in greater depth. The motivation for the two-layer strategy was sparked by our previous observation that DNA loci separated by more than 1.5 Mb on the DNA are likely to be physically located more than 300 nm apart from each other^7, 8^, beyond the diffraction limit. We therefore divided each chromosome into 1.5 Mb blocks and distributed them into three orthogonal fluorescent channels. We then sequentially imaged DNA loci at 25-kb increments in a different block in parallel, circumventing optical crowding, because within the same fluorescent channel, the loci imaged in any given round are genomically at least 4.5 Mb apart from each other. We used 96 rounds of imaging to impart two layer barcodes on 100,049 loci across the genome. The initial 60 rounds resolve 25-kb segments within each chromosome block while the subsequent 36 rounds decode chromosome block identities based on the unique combinations of the pseudocolors across 4 barcoding rounds by leveraging the previous seqFISH+ pseudocolor approach^7, 8, 17^ to the chromosome blocks (Fig. 1b). We have named this imaging-based two-layer barcoding method as two-layer seqFISH+.

We then combined two-layer DNA seqFISH+ with transcriptomic and subnuclear structure measurements to generate multi-omics datasets that integrate information on individual genomic loci (100,049 loci on 20 chromosomes by two-layer DNA seqFISH+), with information on mature and nascent transcripts (up to 1,247 genes based on mRNAs detection by mRNA seqFISH+ or non-barcoded mRNA seqFISH and 17,856 genes based on intron detection by intron seqFISH+) and as well as information on subnuclear structures and chromatin marks based on sequential immunofluorescence assays (using up to 65 antibodies) in thousands of single cells (Fig. 1a-c, Extended Data Fig. 1).

We initially applied this technology to two different mouse cell lines (mouse embryonic stem cells (mESCs), and mammary gland epithelial NMuMG cells), and then compared the results to those from established approaches for benchmarking. We began with two-layer DNA seqFISH+ in mESCs, detecting 63,466 ± 20,525 (median ± s.d.) DNA dots per cell across 100,049 genomic loci in 1,076 cells from two biological replicates (Fig. 1d, top). The estimated detection efficiency was 21.1% while the false positive rate was estimated at 1.2% (Fig. 1d, bottom). We then compared our two-layer DNA seqFISH+ data with previously generated DNA seqFISH+ data^7^ as well as the sequencing-based method, Hi-C^19, 20^, and confirmed that A/B compartments^19, 20^ and other measures are consistent amongst the datasets (Extended Data Fig. 3). Next we examined transcriptomic data from mRNA seqFISH+ (detecting 6,496 ± 2,068 (median ± s.d.) dots per cell), and intron seqFISH+ (1,197 ± 421 (median ± s.d.) transcription active sites per cell) in mESCs, and again showed a high degree of agreement with sequencing-based orthogonal measurements (Extended Data Fig. 2).

**Fig. 2.**
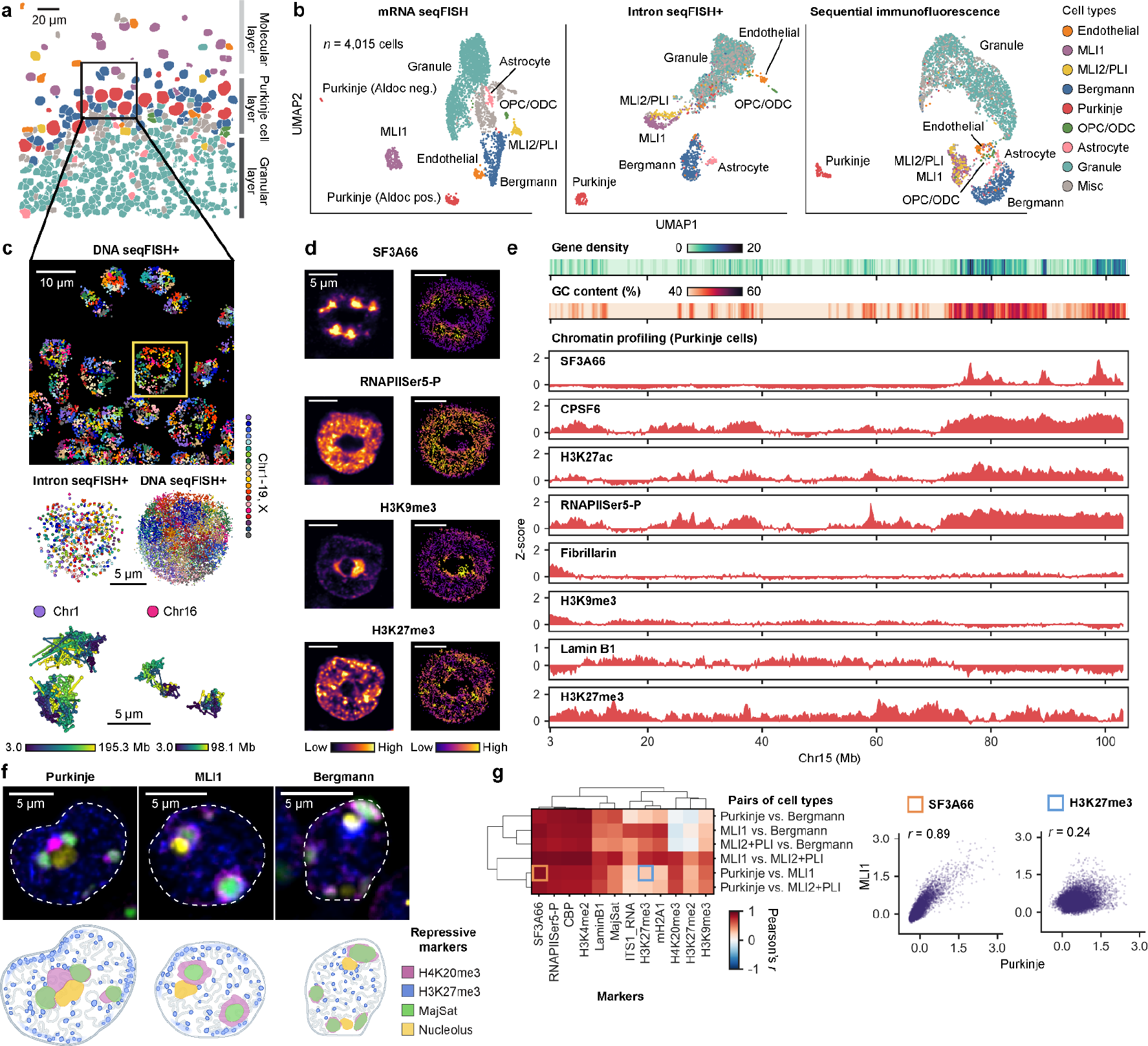
**Single-cell spatial multi-omics in the adult mouse brain cerebellum. a**, Spatial distribution of cell type clusters from a single z-section of the adult mouse cerebellum. **b**, Cell type clusters determined by mRNA seqFISH projected onto UMAP-embedding of mRNA seqFISH (left), intron seqFISH+ (middle), sequential immunofluorescence (right). **c**, Decoded DNA spots (top) from a single z-section of cells in the black box in **a**. Zoomed-in 3D views of decoded intron spots (middle left) and DNA spots (middle right, bottom) in the nucleus from a yellow-boxed Purkinje cell (top). **d**, Immunofluorescence raw images (left) and DNA spots colored by z-score normalized immunofluorescence intensity (right) in the nucleus from a cell highlighted in **c**. **e**, Genomic features and representative imaging-based chromatin profiling in Purkinje cells in chromosome 15. **f**, Representative raw immunofluorescence images from a single z-section for each cell type (top). Illustration showing cell-type specific organization of repressive markers (bottom). The intense H3K27me3 cluster visible in Bergmann glia, representing the inactive X chromosome territory^28^, is not depicted. **g**, Degree of similarity of chromatin profiles between pairs of cell types (left) and corresponding examples (right). *n* = 12,562 loci. 200 kb binning was used for the visualization and analysis (**e**, **g**). *n* = 4,015 cells from two biological replicates of the adult mouse cerebellum in **a**-**g**.

To gain further insight into the spatial organization of these loci, we employed imaging-based chromatin profiling^7, 8^, which measures spatial proximity between genomic loci visualized by two-layer DNA seqFISH+ and subnuclear structures visualized by sequential immunofluorescence (Fig. 1e, Extended Data Fig. 4). We note that to examine subnuclear structures, we included multiple chromatin-associated factors and marks including various histone modifications^21^ (e.g. H3K27ac, H3K9me3) as well as nuclear bodies such as nuclear speckles^22^ (e.g. SF3A66) and nucleolus^23^ (e.g. Fibrillarin), and nuclear lamina^24^ (e.g. Lamin B1). The imaging-based chromatin profiles in mESCs were consistent with those generated by prior sequencing methods as well as previous imaging-based approaches for a wide range of subnuclear structures and histone modifications (e.g. Lamin B1, H3K9ac) (Fig. 1e, f, Extended Data Fig. 4d-h). The imaging-based chromatin profiling spatial resolution is limited by the diffraction-limited immunofluorescence images^25^. Thus, we used 100-200 kb genomic bins, which are typically 100-300 nm away from the adjacent bin for the spatial chromatin profiling analysis.

Nonetheless, because of the improved coverage of genomic loci provided by two-layer DNA seqFISH+, this newly generated chromatin profiling revealed chromatin organization at < 1 Mb resolution (Fig 1e), a level that was inaccessible in previous imaging-based genome-wide studies^5–8^. We will refer to this two-layer DNA seqFISH+ as DNA seqFISH+ for simplicity below. Together, these results confirm the high quality of the spatial multi-omic datasets, which enable us to perform an integrated analysis of nuclear organization across multiple imaging modalities at the submegabase resolution.

### Single-cell spatial multi-omics in the mouse cerebellum

Having established the validity of our approach in mouse cell lines, we proceeded to examine the relationships between chromatin organization and gene regulation in the naive tissue context. We chose the adult mouse cerebellum, a well-defined brain structure with diverse cell types^26^, and examined the cell-type specificity of nuclear organization. Thus we applied DNA seqFISH+ (100,049 loci), intron seqFISH+ (17,856 genes), mRNA seqFISH (60 marker genes), and sequential immunofluorescence (27 markers), to the adult mouse cerebellum.

We identified distinct cell types by mRNA seqFISH transcriptomic profiling and spatially resolved the organization of these diverse cell types in the tissue sections (Fig. 2a-c, Extended Data Figs. 5, 6), capturing the layer organization of the mouse cerebellum at the single-cell resolution. We were able to identify major cell types in the adult mouse brain cerebellum^26^, including neurons such as Purkinje cells, Purkinje layer interneurons (PLI), subtypes of molecular layer interneurons (MLI1 and MLI2), and granule cells, as well as non-neuronal cells such as Bergmann glia, astrocytes, oligodendrocyte precursor cells and oligodendrocytes (OPC/ODC), and endothelial cells. We confirmed the expected layer organization of the adult mouse cerebellum^26^ such as spatially close arrangement of Purkinje cells and Bergmann glia at the Purkinje cell layer, which is adjacent to the granule cell (GC) layer, largely consisting of granule cells (Fig. 2a, Extended Data Fig. 5b, c). Our multi-omics profiling across 4,015 cells further allowed us to compare the genomic landscapes of the different cell types at multiple levels, including nascent transcriptional states by intron seqFISH+ and chromatin states by sequential fluorescence (Fig. 2b, Extended Data Fig. 5). These analyses revealed highly consistent cell-type specific states at each level of analysis (i.e. transcriptional and chromatin).

**Fig. 5.**
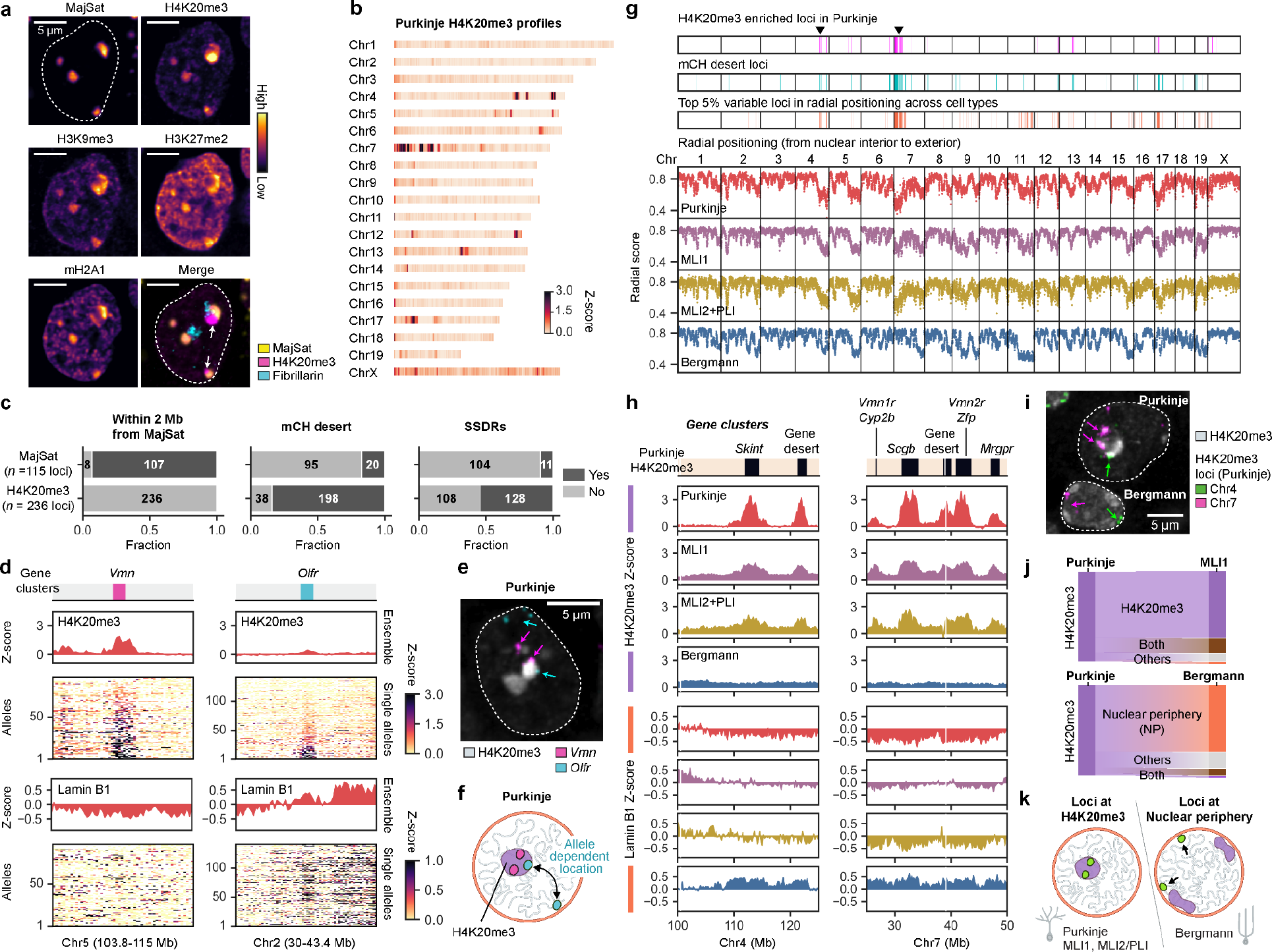
**The H4K20me3 compartment confines specific genomic loci in neurons. a**, DNA FISH (MajSat) and sequential immunofluorescence images for markers enriched near pericentromeric repressive heterochromatin in Purkinje cells. **b**, H4K20me3 enrichment across chromosomes in Purkinje cells. **c**, Barplots comparing the locus characteristics such as mCH desert^48^ and SSDRs^50, 51^ between MajSat- and H4K20me3-enriched loci in Purkinje cells. **d**, Ensemble-averaged and single allele chromatin profiles sorted by H4K20me3 enrichment from bottom to top in Purkinje cells. **e**, Visualization of H4K20me3-enriched *Vmn* and *Olfr* gene family loci with H4K20me3 staining. **f**, Illustration showing the differences of subnuclear localization between *Vmn* (magenta) and *Olfr* (cyan) gene family loci. **g**, Comparison of other genomic features (top) along with radial positioning of chromosomal loci across cell types (bottom). **h**, H4K20me3 and Lamin B1 chromatin profiles at the H4K20me3-enriched regions, highlighted by triangles (**g**). **i**, Visualization of H4K20me3 enriched loci in Chr4 and Chr7 (**g**) overlaid on the H4K20me3 immunofluorescence image in Purkinje and Bergmann cells. **j**, Transition of H4K20me3-enriched loci from Purkinje cells to MLI1 or Bergmann glia. *n* = 252 loci. **k**, Illustration showing the localization switching of genomic loci between neurons and Bergmann glia. 200 kb binning (*n* = 12,562 loci in total) was used for the analysis and visualization. *n* = 128, 263, 88, and 518 cells for Purkinje, MLI1, MLI2+PLI, and Bergmann glia cells from two biological replicates of the adult mouse cerebellum in **a**-**e**, **g**-**j**.

### Cell-type specific active and repressive chromatin compartments in mouse cerebellar cells

Based on the imaging of diverse subnuclear markers together with genome-wide DNA loci by DNA seqFISH+, we were able to identify several different types of chromatin compartments. Active chromatin compartments involved nuclear speckles (SF3A66), which are known to enrich in pre-mRNA splicing factors^22^, and other active chromatin markers such as H3K27ac and RNA polymerase II (RNAPIISer5-P) (Fig. 2d, e, Extended Data Fig. 6d, e). In addition, we also identified at least four major repressive chromatin subcompartments^1, 27^ by examining eight repressive markers, including constitutive heterochromatin (major satellite DNA repeats (MajSat), H4K20me3), facultative heterochromatin (H3K27me3), nucleolus (ITS1 RNA), and nuclear periphery (Lamin B1) (Fig. 2d-f, Extended Data Fig. 6d-g).

**Fig. 6.**
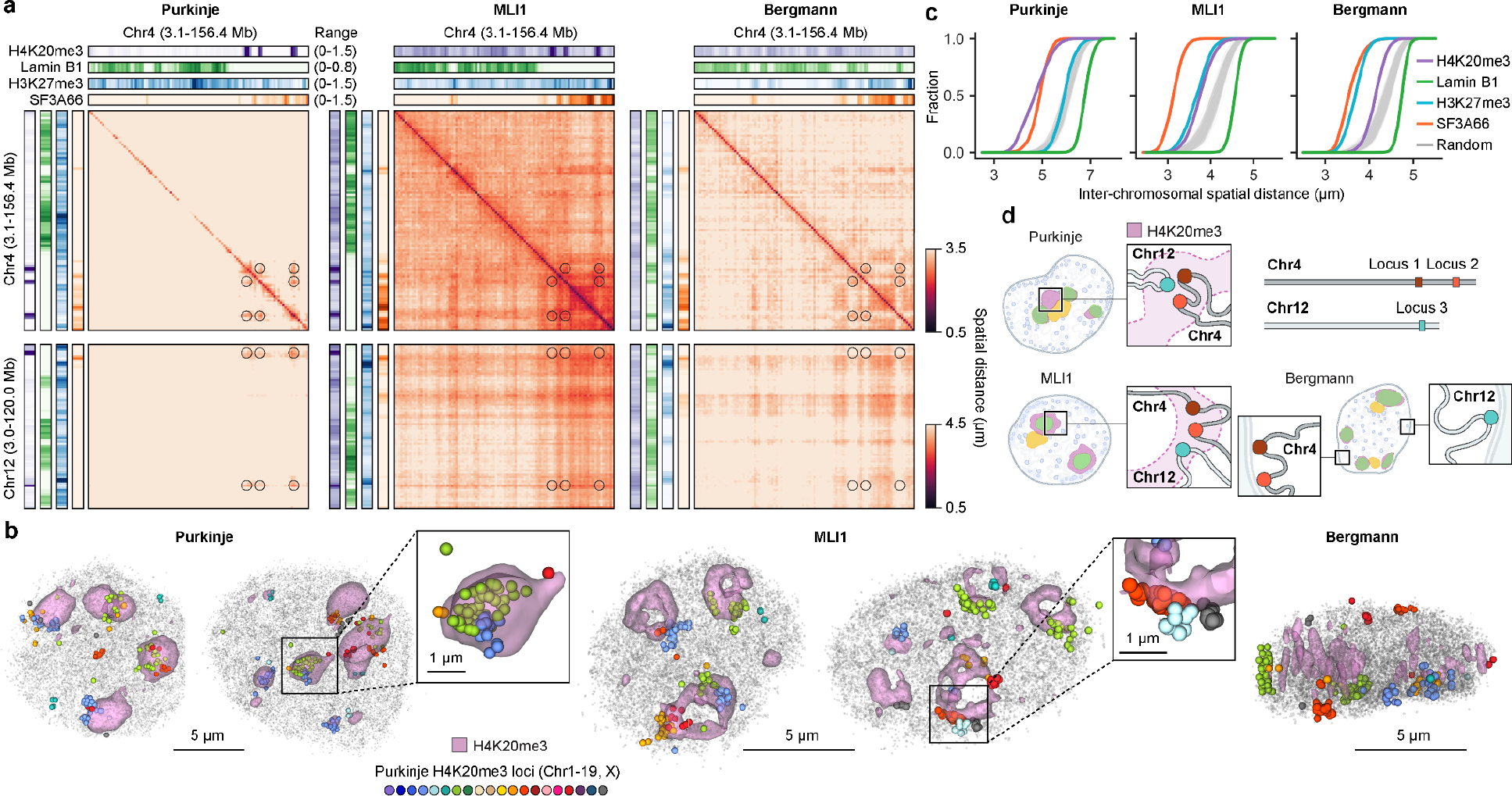
Cell-type specific subnuclear compartmentalization and 3D genome organization. a, Ensemble-averaged spatial distances between pairs of genomic loci along with chromatin profiling at 1.5 Mb resolution from each cell type. Locations of pairs of H4K20me3-associated loci in Purkinje cells in Fig. 5c are highlighted as black circles. **b**, Representative 3D images of H4K20me3 staining and chromosomal loci in each cell type. **c**, Cumulative distribution of inter-chromosomal distances between pairs of loci with top 5% association to a given marker compared to random pairs of loci (*n* = 1,000 trials). 1.5 Mb binning (*n* = 1,678 loci in total), grouped by the chromosome paint block barcodes, was used. **d**, Illustration showing the differences of cell-type specific intra- and inter-chromosomal spatial arrangements around H4K20me3-enriched subnuclear compartment in neurons or at the nuclear periphery in Bergmann glia. *n* = 128, 263, and 518 cells for Purkinje, MLI1, and Bergmann glia cells from two biological replicates of the adult mouse cerebellum in **a**-**c**.

Interestingly, we observed repressive markers were associated with genomic loci in a highly cell-type specific fashion, while active chromatin profiles were largely conserved between cell types at 200 kb resolution (Fig. 2g). For example, by comparing Purkinje and MLI1 cells, we observed that the profiles of active chromatin regions associated with nuclear speckles (marked by SF3A66, as described above) were highly correlated with Pearson correlation coefficient of 0.89, whereas those of repressive regions, marked by H3K27me3, were less so with Pearson correlation coefficient of 0.24 (Fig. 2g, right).

These observations were consistent across various active and repressive markers and in all cell types and cell lines examined (Extended Data Fig. 6h).

Furthermore, some of the repressive markers also showed cell type-specific subnuclear localization patterns. For example, H4K20me3 marker stained subnuclear territories that are segregated from pericentromeric heterochromatin characterized by MajSat^28^ Purkinje cells and MLI1, while those markers stained spatially overlapped regions in Bergmann glia (Fig. 2f, Extended Data Fig. 6f). In addition, regions marked with mH2A1 and H3K27me2 staining, which correlated with H3K27me3 foci in all cell types, also stained pericentromeric regions only in Purkinje cells (Extended Data Fig. 6f). Thus, genomic loci showed cell-type dependent associations with subnuclear compartments of those repressive markers (Extended Data Fig. 6f, g). We examine these active and repressive chromatin compartments and their cell-type specificities in more detail below.

### Active chromatin compartments show distinct sequence features and cell-type specific gene expression patterns

We first focused on transcriptionally active chromatin compartments. Previous work had shown that speckle-associated genomic regions are largely conserved across cell types^8, 29^, but associations with RNAPII have not been fully explored in tissues. In cerebellar cells, genomic regions associated with nuclear speckles tended to be higher in GC content, enriched in shorter-length genes, and also enriched in RNAPIISer5-P marks, regardless of cell type (Fig. 3a-c, Extended Data Fig. 7a, b), consistent with previous works^8, 29–32^. The RNAPIISer5-P broad regions, lacking SF3A66 nuclear speckle marks, formed a distinct compartment far from the speckles (Fig. 3b, Extended Data Fig. 7c), similar to those observed in human cells^30^. Those speckle-enriched and RNAPIISer5-P broad regions not only showed different sequencing features (e.g. GC content) (Fig. 3c, Extended Data Fig. 7b) but also encoded a distinct set of Gene Ontology (GO) terms in each cell type (Extended Data Fig. 7d).

Both nuclear speckles and RNAPIISer5-P broad associated loci showed subtle cell type dependent patterns in their genomic locations (Extended Data Fig. 7e). Nevertheless, we observed that increased association of genomic loci to one of these compartments are typically correlated with cell-type specific gene expression programs (Fig. 3d, e, Extended Data Fig. 7f, g). For example, increased locus associations toward either nuclear speckles or RNAPIISer5-P compartments are coupled with increased gene expression of the loci when comparing Bergmann glia and MLI1 (Fig. 3d, e, bottom). We note that Purkinje cells were an exception, however, where only increased associations to RNAPIISer5-P compartment correlated with increased gene expression (Fig. 3e, Extended Data Fig. 7g).

Furthermore, we classified a third active chromatin compartment associated with RNAPIISer5-P sharp peaks (< 2 Mb), and exhibited a distinct set of characteristics. Genomic loci in this compartment tended to belong to genomic regions with lower GC content, and contained longer genes (Fig. 3c, Extended Data Fig. 7b). In addition, these RNAPIISer5-P sharp loci showed more cell-type specific features, compared to the other active compartments (Extended Data Fig. 7e). This observation motivated us to further characterize the chromatin states and gene expression profiles of long genes (> 200 kb) across cell types. We were particularly interested in these genes, given that neuronal cell types (including cerebellar cells) generally express a greater number of long genes, relative to other cells^33^. In addition, transcriptionally-active long genes have been reported to have unique chromatin signatures *in vivo* including activation-dependent rearrangements^34^, broad enhancer-like chromatin domains^35^, chromatin decondensation^36^, and transcriptional loops^37^.

We found a strong link between the association of long genes with the RNAPIISer5-P subcompartment and their cell-type specific gene expression (Fig. 3f, g). Examples of such genes included Cntnap5b and Dpp10 in Purkinje cells, Pam in MLI1 and MLI2/PLI, Adgrl3 in MLI1 and Bergmann, Slc4a4 in Bergmann, and Kcnd2 and Cadps2 in Granule cells. These long genes tend to play cell-type specific roles in the corresponding cells (e.g. Cadps2, a secretory granule-associated protein and indispensable for normal cerebellar functions^38^), and, indeed, their RNAPIISer5-P enrichment was typically observed only in specific cell types (Fig. 3g). The cell-type specificity was also reflected in the nascent transcription patterns observed by intron seqFISH+ profiles (Fig. 3g) and open chromatin states (Extended Data Fig. 7i). Chromatin profiling, again combining DNA seqFISH+ with sequential immunofluorescence, further identified other subnuclear markers associated with long genes in this compartment, including H3K4me2, H4K8ac, and CBP. However, nuclear speckle marker, SF3A66, typically did not appear at those loci (Fig. 3h, i, Extended Data Fig. 7h), demonstrating a spatial partitioning between nuclear speckles and other active markers (e.g. RNAPIISer5-P, H3K4me2) in single cells (Fig. 3b, Extended Data Fig. 7c). This cell-type specific drastic chromatin reorganization at long gene loci contrasts with the relatively small positional changes observed at gene-dense genomic loci around nuclear speckles^8, 29^. The distinct subnuclear compartments of transcriptionally active loci between nuclear speckles and RNAPII may be functionally important in gene regulation, such as gene expression amplification^32^ and mRNA splicing^39^, and could also be critical in promoting gene misexpression in disease (e.g. speckle-associated loci in schizophrenia^40^).

### Cell-type specific organization of H3K27me3 repressive subnuclear compartments

In contrast to active compartments, genomic loci associated with repressive compartments were highly cell-type specific (Fig 2f, g). We first examined H3K27me3-marked regions, given the role of this repressive compartment in silencing neurodegeneration-related genes^41^. We found there are two major subsets of H3K27me3-associated loci in cerebellar cells. The first set of loci were non-cell-type specific and enriched with genes involved in general developmental processes, represented by gene ontology (GO) terms related to pattern specification, such as the Hox gene clusters on Chr2 and Chr6 (Extended Data Fig. 8a). These associations are consistent with previous data in other biological contexts such as embryonic development^42, 43^. The other set of H3K27me3-associated loci were cell-type specific.

Specifically, Purkinje cells showed an enrichment for the H3K27me3 repressive compartment at genomic loci with longer genes and lower GC content, compared to other cell types such as MLI1 and Bergmann glia (Fig. 4a, Extended Data Fig. 8b). This Purkinje-specific compartment included genes such as Grin2b, whose repression is functionally important in Purkinje cells^44^, and Ptprd, which regulates developmental neurogenesis^45^ (Fig. 4b-d, Extended Data Fig. 8c, d). Some of these long genes (36 out of 46 genes) are developmentally down-regulated from the newborn to the adult Purkinje cells^46^ (Extended Data Fig. 8e).

In addition, we found that H3K27me3- and Lamin B1-enriched loci are negatively correlated in most cell types (Fig. 4b, Extended Data Fig. 8f). However, a subset of H3K27me3-associated loci is enriched with Lamin B1 in Purkinje cells. For example, Ptprd gene locus, spanning for ∼2.2 Mb in Chr4, localized to the nuclear periphery and were marked by Lamin B1 in all major cell types (Purkinje cells, MLI1, and Bergmann glia), but were also marked by H3K27me3 only in Purkinje cells (Fig. 4b-d and Extended Data Fig. 8d). Despite these global organizational differences of the H3K27me3 repressive compartment around the nuclear periphery, the increased association toward either H3K27me3 or Lamin B1 showed decreased nascent transcription levels when comparing the pairs of cell types (Extended Data Fig. 8g). Together, Purkinje cells show a unique organization of their H3K27me3 repressive compartment, perhaps related to their overall highly transcriptionally active nuclei^47^ and increased H3K27me3 modification levels (Extended Data Fig. 5k, l).

### The H4K20me3 subnuclear compartment is associated with specific gene families

We next examined the subnuclear organization of the repressive compartment marked by H4K20me3. Among the pericentromeric repressive markers, H4K20me3 staining marked unique territories adjacent to pericentromeric heterochromatin in Purkinje cells^28^ (Fig. 5a, Extended Data Fig. 9a). We therefore examined the DNA loci associated with H4K20me3 regions in the nucleus to characterize this compartment and their genomic features (Fig. 5b, Extended Data Fig. 9a-c).

We found that H4K20me3-enriched loci in Purkinje cells occur in non-CG methylation (mCH) deserts, i.e. regions that do not accumulate mCH during development^48, 49^, as well as in mouse strain-specific diverse regions (SSDRs)^50, 51^ (Fig. 5c, Extended Data Fig. 9d). Those enrichments contrasted from spatially adjacent pericentromeric MajSat-associated loci, which are mostly found within 2 Mb from chromosome start coordinates (Fig. 5c, Extended Data Fig. 9a). Moreover, the H4K20me3-enriched loci appeared on specific chromosomes, including Chr7 and Chr17 (Fig. 5b) and included both gene-coding regions and gene deserts. In the gene-coding regions, we identified gene clusters, such as vomeronasal receptors (*Vmn*), secretoglobins (*Scgb*), and zinc finger proteins (*Zfp*) (Extended Data Fig. 9e), some of which (e.g. *Vmn*, *Zfp*) are distributed across multiple chromosomes. We also found that *Vmn* gene clusters are consistently marked by H4K20me3, and do not show nuclear lamina association (Fig. 5d-f,

Extended Data Fig. 9h, i). In contrast, other gene family clusters such as olfactory receptors (*Olfr*) had a more mixed profile, associating with H4K20me3 in some cells and some alleles while with Lamin B1 at the nuclear periphery in others. This feature gave *Olfr* genes overall lower H4K20me3 enrichments in Purkinje cells (Fig. 5d-f, Extended Data Fig. 9h, i), despite showing similar sequencing features (i.e. mCH deserts and SSDRs) as *Vmn* family genes^48, 50, 51^. Taken together, these data reveal that H4K20me3-marked regions constitute a separate subnuclear compartment with highly specific locus associations.

### H4K20me3 supports cell-type specific radial chromatin organization

We noticed that certain chromosomal loci such as those in Chr4, Chr7, and Chr17 showed neuron-specific interior radial positioning, different from their arrangement in Bergmann glia (Fig. 5g, Extended Data Fig. 9j). Given that a majority of these loci were enriched for H4K20me3 in Purkinje cells, we wondered whether H4K20me3 may be related to the cell-type specific radial positioning of chromosomes. Consistent with this notion, we found that H4K20me3 territories tend to form in the nuclear interior in Purkinje cells^52^ (Fig. 5a), and H4K20me3-associated loci were therefore also localized to the interior (Fig. 5g, Extended Data Fig. 9k). For example, H4K20me3-associated *Vmn* gene clusters are found at 2.7 ± 0.6 μm (median ± s.d.) interior from the nuclear periphery, in contrast to 1.5 ± 0.5μm (median ± s.d.) for the weakly H4K20me3-associated *Olfr* gene clusters and 1.3 ± 0.5 μm (median ± s.d.) from all genomic loci in Purkinje cells (Extended Data Fig. 9e).

Interestingly, we found the loci enriched in H4K20me3 in Purkinje cells were highly conserved with other neurons in their association with the H4K20me3 subcompartment (Fig. 5h-j, Extended Data Fig. 9l, m; 90.5% with MLI1 and 92.5% with MLI2/PLI). However, those loci were not enriched with H4K20me3 in Bergmann glia (8.7%), where they localized at the nuclear periphery and showed enrichment for Lamin B1 (Fig. 5h-j, Extended Data Fig. 9m). For example, the *Skint* gene cluster and the gene desert regions, which are ∼6.8 Mb apart in Chr4, were enriched with H4K20me3 at the nuclear interior and depleted for Lamin B1 in three types of neurons, while the same genomic regions were not marked by H4K20me3 and instead showed enrichment with Lamin B1 and localization to the nuclear periphery in Bergmann glia (Fig. 5g-k). By contrast, chromosomal loci in Chr11 and Chr19 showed more interior radial positioning in glial nuclei compared to neurons (Fig. 5g). We had observed similar differences between neuron versus glia radial chromosomal positioning in the adult mouse cerebral cortex^8^, where Chr7 and Chr17 are positioned in the nuclear interior in neurons whereas Chr11 and Chr19 are interior in astrocytes. These results suggest that different brain regions show conserved nuclear organization patterns that are distinct between neurons and glial cells.

### Subnuclear chromatin compartments underpin the 3D organization of the genome

Having examined the features of each separate chromatin compartment, we next investigated the spatial relationship of the genome organization with the subnuclear structures in the different cell types of the adult mouse cerebellum. To do so, we systematically calculated the average inter-chromosomal distances between pairs of genomic loci enriched with subnuclear markers using top 5% genomic loci associated with each marker (Fig. 6, Extended Data Fig. 10). We observed that, in all cell types, pairs of genomic loci enriched with the same specific markers - such as markers for nuclear speckles and pericentromeric heterochromatin - show closer average inter-chromosomal distances compared to those with random selection (Fig. 6a, c, Extended Data Fig. 10a-c), consistent with previous literature^7, 8, 31, 53–56^. Lamin B1-enriched loci were an exception, showing longer spatial distances between chromosome pairs relative to controls, suggesting a lack of interaction (Fig. 6a, c, Extended Data Fig. 10a, b). At the other end of the spectrum, H3K27me3-enriched loci were on average closer to each other than random pairs in MLI1 and Bergmann glia but not in Purkinje cells. These observations can be explained by the fact that loci localizing to the nuclear periphery, which generally encompasses a larger area, are more likely to be distant from each other, than pairs of loci in the nuclear interior. Thus, pairs of chromosomes with exterior radial positioning at the nuclear periphery tend to be spatially farther away from each other.

Consistent with this notion, chromatin profiles showed that H3K27me3-enriched loci localized to the interior in MLI1 and in Bergmann glia, but appear at the nuclear periphery in Purkinje cells, while Lamin B1-enriched loci are at the nuclear periphery in all cell types (Extended Data Fig. 8f).

We further examined the relationships between subnuclear compartmentalization and 3D genome organization using the H4K20me3-enriched loci identified in Purkinje cells (Figs. 5g, 6b, Extended Data Fig. 10c, d). Consistent with the above model, loci enriched in H4K20me3, which are generally located at the nuclear interior in neurons such as Purkinje and MLI1, exhibited increased inter-chromosome association (i.e. shorter distances) (Fig. 6b, d, Extended Data Fig. 10c, d). In contrast, in Bergmann glia, where those H4K20me3 loci are present at the nuclear periphery (Fig. 5j), we observed decreased inter-chromosomal association of the loci (i.e. increased distances) (Fig. 6a, Extended Data Fig. 10c). However, we note long-range intra-chromosomal associations between the Purkinje H4K20me3-enriched loci were conserved in all three cell types (Fig. 6a, Extended Data Fig. 10c). The differences in inter-chromosomal versus intra-chromosomal associations are likely accounted for by the geometric larger area of nuclear periphery versus nuclear interior, which may also explain the lack of inter-chromosomal interactions in lamin associated loci (Fig. 6d). Together, these data support a model in which subnuclear compartments help to shape the 3D genome organization in the nucleus in a cell-type specific fashion.

## Discussion

Here we demonstrated how high-resolution seqFISH-based single-cell multi-omics profiling can reveal cell-type specific chromatin compartmentalization in native tissues, and identify specific genes associated with each compartment. We showed that our two-layer barcoding strategy can effectively cover a large number (>100,000) of genomic loci across the genome, larger than what is feasible with existing imaging-based omics methods^16–18^. This large-scale barcoding capability enables comprehensive *in situ* investigation of a diverse set of RNAs, including splicing variants and non-coding RNAs^57, 58^, as well as chromosome abnormalities at the submegabase resolution in a variety of tissues, including tumors^59^.

The high-resolution spatial multi-omics data revealed that repressive chromatin compartments are globally organized in a more cell-type specific fashion compared to active chromatin compartments in the adult mouse cerebellum, while subtle changes of genomic loci toward active subnuclear compartments between cell types are tightly linked to cell-type specific gene expression programs. We note that the resolution of the current analysis is limited with diffraction-limited immunofluorescence images and can be further increased with super-resolution imaging of the chromatin marks^25^.

The cell-type specific repressive and active subnuclear compartments are correlated with variations in the chromosomal structures in different cell types. In particular, we demonstrated that a H4K20me3 compartment, localized adjacent to pericentromeric heterochromatin^28^, confines genomic loci enriched with a subset of specific gene families (e.g. *Vmn*, *Zfp*) with a high specificity. Within the adult mouse cerebellum, this confinement was characterized at the nuclear interior in neuronal cells, but not Bergmann glia. In contrast, the same set of loci switched to nuclear lamina association at the nuclear periphery in Bergmann glia cells. Thus, those genomic loci contribute to cell-type specific 3D genome organization, including radial positioning of chromosomes, as well as intra- and inter-chromosomal interactions. Previously, H4K20me3 compartments had been shown to be involved in chromosomal organization of olfactory receptor genes in the olfactory sensory neurons^54, 60–63^. Our results now reveal a broader role for H4K20me3 in organizing the 3D genome with a different set of genomic loci (e.g. *Vmn*, *Skint* gene clusters), in a cell-type-dependent manner. Finally, the H4K20me3 compartmentalization in neurons may account for a wide-range of 3D genome organization, including the neuron-specific radial reorganization of chromosomes during postnatal development^49^ and long-range intra- and inter-chromosomal interactions in the other mouse brain regions^36, 49^. The H4K20me3 compartment could also be disrupted in neurological and neurodegenerative disorders^28, 64, 65^ and can be investigated further with the seqFISH-based high-resolution spatial multi-omics approach.

## Acknowledgements

We thank I. Strazhnik for help with figures; I. Carmi, P. Bhat, and M. Elowitz for help with manuscript; M. Guttman, C. H. Tischbirek, S. Shah, and C.-H. L. Eng for helpful discussion; C. Karp, C. Cronin and A. Cronin for help with automated imaging setup; F. Gao (Bioinformatics Resource Center in the Beckman Institute at Caltech) for processing published sequencing data; S. Peiró and M. A. Marti-Renom for sharing processed Hi-C data in NMuMG cells; N. J. Rezaee and A. Cunha for help with cell segmentation; This project is funded by NIH U01DK127420 and Allen Discovery Center.

## Author contributions

Y.T. and L.C. conceived the idea and designed experiments. Y.T., J.Y., and S.S. prepared and validated experimental materials. Y.T. performed imaging experiments. Y.T. and J.W. performed image processing with help from L.J.O. Y.T. and Y.Y. performed data analysis with help from M.P. Y.T. and L.C. wrote the manuscript with input from all authors. L.C. supervised the project.

## Materials & Correspondence

should be addressed to Yodai Takei or Long Cai.

## Data availability

The source data from this study are available at Zenodo (https://zenodo.org/record/7693825; doi: 10.5281/zenodo.7693825). Additional processed data and experimental resources (e.g. probe sequences) from this study are available at GitHub (https://github.com/CaiGroup/dna-seqfish-plus-multi-omics). The raw microscopy data are not uploaded owing to their large size (6.8 Tb), but are available upon reasonable request. Publicly available datasets used in this study (GSE98674, GSE96033, GSE48895, GSE166210, GSE181693, GSE17051, GSE165371, 4DNESL2AY9CM) are detailed in the Methods. The chromosomal DNA binding probe sequences were obtained from mm10 newBalance DNA FISH probes at PaintSHOP resources (https://github.com/beliveau-lab/PaintSHOP_resources).

## Code availability

The custom written scripts used in this study are available at GitHub (https://github.com/CaiGroup/dna-seqfish-plus-multi-omics).

## Methods

### Data reporting

No statistical methods were used to predetermine sample size. The experiments were not randomized, and the investigators were not blinded to allocation during experiments and outcome assessment.

### RNA seqFISH+ encoding strategy

To spatially resolve mRNA profiles for 1,194 genes in the cell culture experiments (see ‘Cell culture experiment’), we used a modified version of RNA seqFISH+ encoding scheme^17^ consisting of 16-pseudocolor bases with 4 rounds of barcoding including one-error correction round, which can accommodate up to 4,096 (= 16^3^) genes, in one fluorescent channel (635 nm). Additional 64 genes for mRNA and non-coding RNA species were encoded as a non-barcoded seqFISH scheme^7, 8, 16, 78^ in one fluorescent channel (488 nm). In addition, to spatially resolve the nascent transcriptome (17,856 genes) typically at their transcription active sites in the nucleus^16^, we applied a modified version of RNA seqFISH+ encoding scheme^17^ consisting of 12-pseudocolor bases with 5 rounds of barcoding including one-error correction round, which can accommodate up to 20,736 (= 12^4^) genes, in one fluorescent channel (561 nm). This intron seqFISH+ approach allows us to super-resolve subcellular localization of intronic RNAs at the sub-pixel resolution while original intron seqFISH decoding for 10,421 genes^16^ was performed with pixel resolution using three fluorescent channels.

Similarly, for mouse brain cerebellum experiments (see ‘Tissue slice experiment’), we profiled 60 mRNA species including cell-type marker genes in the adult mouse brain cerebellum^26^ with the non-barcoded seqFISH strategy in one fluorescent channel (635 nm) as well as 17,856 genes for nascent transcripts by intron seqFISH+ in one fluorescent channel (561 nm) as described above.

### Two-layer DNA seqFISH+ encoding strategy

To spatially resolve whole mouse chromosomes with 25-kb resolution, a two-layer barcoding strategy, consisting of diffraction limited locus imaging and chromosome paint imaging, was developed. First, mm10 mouse genome was divided into non-overlapping 25-kb loci and up to 60 loci were grouped together to make 1.5-Mb chromosome paint blocks. Those chromosome paint blocks were then separated into three groups according to their genomic coordinates in order to be encoded by three orthogonal fluorescent channels (*n* = 560, 559, and 559 blocks used in 635 nm, 561 nm, and 488 nm fluorescent channels). In total, 96 rounds of imaging were performed to decode the 100,049 loci encoded in two-layer DNA seqFISH+ (Supplementary Table 1). In the initial 60 rounds of imaging, 25-kb loci were sequentially read out one at a time for all chromosome paint blocks based on their genomic coordinates within each block in each fluorescent channel. These 60 rounds can resolve the identities of 25-kb loci within each chromosome paint block but cannot distinguish which specific chromosome paint block those loci belong to. In the subsequent 36 rounds of imaging, chromosome paint block identities were decoded by painting the individual 1.5-Mb blocks using a 9-pseudocolor base seqFISH+ coding scheme^17^ with 4 rounds of barcoding in each fluorescent channel. This allows to resolve up to 729 (= 9^3^) chromosome paint blocks in each fluorescent channel with one extra round for a stringent decoding.

While original implementations of seqFISH+^7, 8, 17^ barcoded individual diffraction limited spots, this strategy barcodes individual chromosome paint blocks with unique pseudocolor combinations. This two-layer DNA seqFISH+ strategy, leveraging two layers of orthogonal barcoding (i.e. sequential barcoding of diffraction limited loci and scalable barcoding of chromosome paint blocks), can efficiently encode up to 131,220 (= 60 × 729 × 3) genomic loci within 96 rounds in three fluorescent channels, which are sufficient to accommodate all 25-kb loci in the mouse and human genome.

### Primary probe design and synthesis

mRNA seqFISH+, intron seqFISH+, and non-barcoded RNA seqFISH probes were designed as described before^7, 8, 16, 17^. In brief, 35-nt RNA target binding sequences, 15-nt readout probe binding sites^7, 8^, and a pair of 20-nt primer binding sites at 5’ and 3’ end of the primary probes were concatenated (up to 150-nt for mRNA seqFISH+ and non-barcoded RNA seqFISH; up to 170-nt for intron seqFISH+) to allow enzymatic probe amplification steps below. We used 32 primary probes per gene for mRNA seqFISH+, 12-25 primary probes (25 probes whenever possible) per gene for intron seqFISH+, and 8-50 primary probes per gene for non-barcoded RNA seqFISH.

For two-layer DNA seqFISH+ primary probes, the chromosomal DNA binding sequences were selected from mm10 newBalance DNA FISH probes at PaintSHOP resources^79^(https://github.com/beliveau-lab/PaintSHOP_resources). Specifically, primary probes with reported off-target scores less than or equal to 200 were initially selected and the total number of probes in each defined 25-kb locus was counted. Then the primary probes were sorted by the off-target score and 30 probes were selected from the smallest off-target score per genomic locus while the genomic locus with less than 30 probes were filtered out. After the selection, the primary probe sequences were assembled with readout probe and primer binding sites similarly to the DNA seqFISH+ study^7, 8^ with modified combinations of readout probe binding sites based on the two-layer DNA seqFISH+ coding scheme (see ‘Two-layer DNA seqFISH+ encoding strategy’). At each 25-kb locus targeted, we used 30 primary probes selected with the criteria above to image individual loci as diffraction limited spots. Those primary probes (up to 170-nt) consist of the genomic region specific sequences (30-37-nt)^79^ flanked by spacer sequences (“AA” and “A”), three identical 15-nt readout binding sites, corresponding to one of the 60 rounds of sequential diffraction limited spot imaging (hybridizations 1-60) in each fluorescent channel, three 15-nt readout binding sites, corresponding to three out of four chromosome paint barcoding rounds (hybridizations 61-96) in each fluorescent channel, and a pair of 20-nt primer binding sites at 5’ and 3’ end of the primary probes.

RNA seqFISH+ and two-layer DNA seqFISH+ primary probes were generated from oligo array pools (Twist Bioscience for 150-nt oligos; Agilent Technologies for 170-nt oligos) based on Oligopaint technologies^80^ with enzymatic amplifications as described previously^7, 8, 16, 17^. Total of 3,001,470 two-layer DNA seqFISH+ primary probes were obtained from 13 different oligo array pools. Briefly, we performed PCR with Q5 Hot Start High-Fidelity (NEB M0494S), column purification with QIAquick PCR Purification Kit (Qiagen 28104), *in vitro* transcription (NEB E2040S) at 42°C for 8 hours in the presence of RNasin Ribonuclease Inhibitor (Promega N2111) and Pyrophosphatase (NEB M0361S), and reverse transcription (Thermo Scientific EP0751) at 50°C for 2 hours and then at 55°C for 2 hours followed by heat inactivation at 85°C for 5 minutes, RNA hydrolysis by 1 M NaOH at 65°C for 15 minutes, and neutralization by an equal amount of 1 M acetic acid. Then generated primary probes were concentrated by ethanol precipitation, pooled if necessary, then dried to a powder by speed-vac, and stored at -20°C. The primer pairs for the amplification were selected from previous studies^7, 8^.

The DNA FISH probes for mouse repetitive elements (LINE1, SINEB1, Telomere, MinSat, MajSat, and rDNA) and 3632454L22Rik fiducial marker were designed and synthesized as described before^7, 8^.

Similarly, RNA FISH probes for mouse repetitive elements were designed and synthesized for this study. In brief, SINEB1 and two orthogonal sets of MERVL RNA FISH probes were generated using genomic DNA of E14 mESCs extracted with DNeasy Blood and Tissue Kits (Qiagen 69504) as a template for PCR, followed by in vitro transcription and reverse transcription. The PCR primers consisted of genomic DNA binding sites, originally designed for RT-qPCR^81^, and overhangs of readout probe binding sites. Single RNA FISH probes targeting MajSat (both sense and antisense strands), telomeric repeat-containing RNA (TERRA), and Nsmce2 intronic RNA repetitive regions used as internal fiducial markers were designed with overhangs of readout probe binding sites and purchased from Integrated DNA Technologies.

### Readout probe design and synthesis

To implement the two-layer DNA seqFISH+ strategy, 96 unique readout probes were used in each fluorescent channel for a total of 288 unique readout probes for 3 fluorescent channels. The readouts probe sequences were obtained from our previous DNA seqFISH+ studies^7, 8^ as well as additional orthogonal readout probe sequences were generated and validated similarly to those previous studies. The RNA seqFISH+ readout probe sequences were selected from a subset of the two-layer DNA seqFISH+ readout probe sequences. The readout probe sequences (12-15-nt) for sequential immunofluorescence were selected from our previous studies^7, 8^ and further designed and validated with the same criteria for this study. The fluorescently-labeled readout probes (Integrated DNA Technologies) that can bind to the readout sequences on the primary probes or primary antibodies were conjugated in-house to Alexa Fluor 647–NHS ester (Invitrogen A20006), Cy3B–NHS ester (GE Healthcare PA63101), or Alexa Fluor 488–NHS ester (Invitrogen A20000) as described before^17^ or directly purchased (Integrated DNA Technologies).

### DNA**-**antibody conjugation

The oligonucleotide-conjugated antibodies were prepared similarly to those previously described^7, 82, 83^. The BSA-free primary antibodies were purchased from commercial vendors whenever possible. For the BSA-free primary antibodies, we used the crosslinking of 5’ thiol-modified 18-nt DNA oligonucleotides (Integrated DNA Technologies) to lysine residues on antibodies via PEGylated SMCC cross-linker (SM(PEG)2) (Thermo Scientific Thermo Scientific 22102) ^7, 83^. In addition, for some of the BSA-free primary antibodies, we used the crosslinking of 5’ azide-modified 18-nt DNA oligonucleotides (Integrated DNA Technologies) to lysine residues on antibodies via DBCO-PEG4-NHS (Sigma-Aldrich 764019) cross-linker with modifications from previous protocols^82^. In brief, primary antibodies (90-100 μg) were buffer-exchanged to 1× PBS (Invitrogen AM9624) using 50KDa Amicon Ultra Centrifugal Filter Unit (Millipore, UFC505096) and reacted with 10 equivalents of DBCO-PEG4-NHS at 4°C for 4–6 hours. Then the antibody solution was exchanged with 1× PBS using the 50KDa Amicon Ultra Centrifugal Filter Unit and reacted with 10 equivalents of the azide-modified 18-nt DNA oligonucleotides at 4°C for 48 hours. The DNA-primary antibody conjugates were washed with 1× PBS and concentrated using the 50KDa Amicon Ultra Centrifugal Filter Unit. For BSA-containing primary antibodies, we used SiteClick R-PE Antibody Labeling Kit (Life Technologies S10467) to crosslink 5’ DBCO-modified 18-nt DNA oligonucleotides (Integrated DNA Technologies) to the specific sites on primary antibodies^7, 83^. In all conjugation strategies, excess oligonucleotides in the antibody solution were removed using the 50KDa Amicon Ultra Centrifugal Filter Unit at the final step. The oligonucleotide-conjugated primary antibodies were individually validated by immunofluorescence for their subcellular localization patterns and stored in 1× PBS at −80 °C as small aliquots. The oligonucleotide-conjugated primary antibodies are listed in Supplementary Table 2.

### Cell culture and preparation

E14 mouse embryonic stem cells (E14Tg2a.4) (RRID:MMRRC_015890-UCD) from Mutant Mouse Regional Resource Centers were maintained on 0.1% gelatin (Sigma-Aldrich G1393) coated 6-well plates under the serum/LIF condition containing 15% ES-grade FBS (Gibco 16141061), 1,000 units/mL leukemia inhibitory factor (LIF) (Sigma-Aldrich ESG1106), 1× non-essential amino acids (Gibco 11140050), 1 mM sodium pyruvate (Gibco 11360070), 55 μM 2-mercaptoethanol (Gibco 21985023), 1× penicillin and streptomycin (Gibco 15140122) in DMEM GlutaMAX (Gibco 10566016). NMuMG mammary gland cells (ATCC CRL-1636) were maintained on 6-well plates (Thermo Scientific 150687) in 10% FBS (Corning 35-010-CV), 10 μg/mL insulin (Gibco 12585014), 1× penicillin and streptomycin (Gibco 15140122), and DMEM (Corning 10-013-CV).

The coverslips were prepared as previously described^7^. In brief, E14 cells were plated on poly-D-lysine (Sigma-Aldrich P6407) and human laminin (BioLamina LN511) coated coverslips (25 mm x 60 mm) and incubated for 24 hours. Similarly, NMuMG cells were plated on poly-D-lysine (Sigma-Aldrich P7280) and human laminin (BioLamina LN511) coated coverslips and incubated for 24 hours. The cells were washed with 1× PBS (Invitrogen AM9624) once and fixed with freshly prepared 4% formaldehyde (Thermo Scientific 28908) in 1× PBS at room temperature for 10 minutes. The fixed cells were washed with 1× PBS a few times and stored in 70% ethanol at -20°C^7, 84^ until the cell culture experiment below.

### Cell culture experiment

The samples for cell culture experiments were prepared similarly to our previous study^7^ with some modifications. First, the samples were permeabilized and prepared for sequential immunofluorescence. The fixed cells on the coverslips were dried and permeabilized with 0.5% Triton-X (Sigma-Aldrich 93443) in 1× PBS at room temperature for 15 minutes using a sterilized custom-made chamber with a silicon plate (McMASTER-CARR 86915K16) attached on each coverslip. The samples were then washed three times with 1× PBS and blocked at room temperature for 15 minutes with a blocking solution consisting of 1× PBS, 10 mg/mL UltraPure BSA (Invitrogen AM2616), 0.3% Triton-X, 0.1% dextran sulfate (Sigma D4911) and 0.5 mg/mL sheared Salmon Sperm DNA (Invitrogen AM9680). Then DNA oligo-conjugated primary antibodies were incubated in the blocking solution with 100-fold diluted SUPERase In RNase Inhibitor (Invitrogen AM2694) at 4°C overnight. The typical estimate of the final concentration of each primary antibody in the blocking solution was 5-10 ng/μL. After DNA oligo-conjugated primary antibody incubation, the samples were washed with 1× PBS three times and incubated at room temperature for 15 minutes. The samples were then post-fixed with freshly made 4% formaldehyde in 1× PBS at room temperature for 5 minutes, washed with 1× PBS six times, and incubated at room temperature for 15 minutes. The samples were further post-fixed with 1.5 mM BS(PEG)5 (PEGylated bis(sulfosuccinimidyl)suberate) (Thermo Scientific A35396) in 1× PBS at room temperature for 20 minutes and quenched with 100 mM Tris-HCl pH7.4 (Alfa Aesar J62848) at room temperature for 5 minutes. We note these post-fixation steps allow the stabilization of antibodies and samples during the heating step required for DNA FISH preparation. After the post-fixation steps, the samples were washed with 1xPBS three times and air dried upon removing the custom silicon chamber.

After the sequential immunofluorescence preparation above, the samples were prepared for RNA seqFISH steps. The custom-made flow cells (fluidic volume ∼30 μl), made from glass slide (25 x 75 mm) with 1 mm thickness and 1 mm diameter holes and a PET film coated on both sides with an acrylic adhesive with total thickness 0.25 mm (Grace Bio-Labs RD481902), were attached to the prepared coverslips. The samples were rinsed with 4× SSC and hybridized with RNA seqFISH primary probe pools (1-10 nM per probe) and 10 nM polyT LNA oligo (Qiagen) in a 50% hybridization buffer consisting of 50% formamide (Invitrogen AM9342), 2× SSC, and 10% (w/v) dextran sulfate (Millipore 3710-OP). The hybridization was performed at 37°C for 48-72 hours in a humid chamber. After primary probe hybridization, the samples were washed with a 55% wash buffer consisting of 55% formamide, 2× SSC, and 0.1% Triton X-100 at room temperature for 30 minutes and rinsed three times with 4× SSC. Then the samples were imaged for RNA seqFISH as described below (see ‘Sequential imaging’).

After RNA seqFISH imaging, the samples were prepared for DNA seqFISH+ steps. The samples were taken from the microscope, rinsed with 1× PBS, and incubated with 100-fold diluted RNase A/T1 Mix (Thermo Fisher EN0551) in 1× PBS at 37°C for 1 hour to digest RNA species and remove RNA seqFISH+ primary probes. The samples were then rinsed three times with 1× PBS and three times with a 50% denaturation buffer consisting of 50% formamide and 2× SSC and incubated at room temperature for 15 minutes. Following the incubation, the samples were heated on the heat block at 90°C for 5 minutes in the 50% denaturation buffer with aluminum sealing tapes (Thermo Scientific 232698) on the inlet and outlet of the custom chamber. Immediately after heating, the samples were rinsed with 4× SSC and hybridized with a DNA seqFISH+ primary hybridization buffer consisting of two-layer DNA seqFISH+ probes (∼0.2 nM per probe), rDNA probes (∼10 nM per probe), ∼1 μM LINE1 probe, ∼1 μM SINEB1 probe, 100 nM 3632454L22Rik fiducial marker probe (Integrated DNA Technologies), 35% formamide, 2× SSC, and 10% (w/v) dextran sulfate (Millipore 3710-OP) at 37°C for 5-7 days in a humid chamber. After the hybridization step, the samples were washed with a 35% wash buffer, consisting of 35% formamide, 2× SSC and 0.1% Triton X-100, for six times and then incubated at room temperature for 15 minutes, followed by rinsing for three times with 4× SSC.

After DNA seqFISH+ hybridization, the samples were further processed to stably maintain DNA seqFISH+ primary probes on the chromosomal DNA during imaging routines. First, 250 nM global ligation bridge oligo^7, 8^(Integrated DNA Technologies) was hybridized in a 20% hybridization buffer consisting of 20% formamide, dextran sulfate (Sigma D4911), and 4xSSC at 37°C for 2 hours. The samples were then washed three times with a 12.5% wash buffer consisting of 12.5% formamide, 2× SSC, and 0.1% Triton X-100, incubated at room temperature for 5 minutes, and rinsed three times with 1× PBS. Next, to perform ligation between 5’- and 3’-end of the DNA seqFISH+ primary probes with the hybridized ligation bridge oligo, the samples were incubated with 20-fold diluted Quick Ligase in 1× Quick Ligase Reaction Buffer from Quick Ligation Kit (NEB M2200) supplemented with additional 1 mM ATP (NEB P0756) at room temperature for 1 hour. The samples were then washed with the 12.5% wash buffer and rinsed three times with 1× PBS. After the ligation steps above, the samples were further processed with amine modification and post-fixation to synergistically stabilize the probes as we demonstrated before^7^. For the amine modification, the samples were rinsed with 1× Labeling Buffer A and incubated with 10-fold diluted Label IT Amine Modifying Reagent in 1× Labeling Buffer A from Label IT Nucleic Acid

Modifying Reagent (Mirus Bio MIR 3900) at room temperature for 45 minutes. After three rinses with 1× PBS, the samples were post-fixed with 1.5 mM BS(PEG)5 in 1× PBS at room temperature for 30 minutes and quenched with 100 mM Tris-HCl pH7.4 at room temperature for 5 minutes. Then the samples were washed with the 55% wash buffer at room temperature for 5 minutes, rinsed with 4× SSC for three times, and stored in 4× SSC at 4°C until the imaging for DNA seqFISH+ and sequential immunofluorescence (see ‘Sequential imaging’).

### Tissue slice experiment

All animal care and experiments were carried out in accordance with Caltech Institutional Animal Care and Use Committee (IACUC) and NIH guidelines. 6-week-old C57BL/6J female mice from The Jackson Laboratory (Stock No: 000664 | B6) were used for the cerebellum tissue slice experiments. The brain samples and coverslips with 15-20 μm coronal sections of the cerebellum were prepared similarly to those described before^8^.

The tissue slice experiments were performed similarly to the cell culture experiment (see ‘Cell culture experiment’) and our previous mouse cortex experiment^8^ with some modifications. In brief, the permeabilization and sequential immunofluorescence were performed as described before^8^. After the sequential immunofluorescence preparation, custom-made flow cells (fluidic volume about 40 μl) were attached to the coverslips. Then the RNA seqFISH preparation and imaging were performed similarly to the cell culture experiment with a different set of non-barcoded mRNA seqFISH primary probes including mouse cerebellum marker genes^26^, intron seqFISH+ probes, and polyT LNA oligo (Qiagen). After RNA seqFISH imaging, the samples were prepared for DNA seqFISH+ steps similarly to the cell culture experiment except the extended 90°C heating time to 6 minutes, followed by DNA seqFISH+ and sequential immunofluorescence imaging as described below (see ‘Sequential imaging’).

### Automated microscope setup

All imaging experiments were performed with the confocal fluorescence imaging platform and fluidics delivery system as described before^7, 8, 16, 17^. In brief, the microscope (Leica DMi8) was equipped with a confocal scanner unit (Yokogawa CSU-W1), a sCMOS camera (Andor Zyla 4.2 Plus), a 63× oil objective (NA

= 1.40, Leica 11506349), Borealis beam conditioning unit (Andor), a motorized stage (ASI MS2000), fiber coupled lasers (635, 561, 488, and 405 nm) from CNI and Shanghai Dream Lasers Technology, and filter sets from Semrock. In addition, the custom-made automated sampler was set up for automated buffer exchange coupled with hybridization and imaging routines (see ‘Sequential imaging’) to move to the well of the designated hybridization buffer corresponding to each hybridization round from a 2.0-mL 96-well plate (Corning 3960). The hybridization buffer and other buffers were moved through a multichannel fluidic valve (IDEX Health & Science EZ1213-820-4) to the custom-made flow cell with a syringe pump (Hamilton Company 63133-01). The automated fluidics delivery and imaging were controlled by a custom-written script in μManager^85^.

### Sequential imaging

The seqFISH hybridization and imaging routines were performed as described previously^7, 8, 16, 17^. In brief, the sample with the custom-made flow cell was connected to the automated fluidics system on the microscope. The field of views (FOVs) for the images were registered based on the DAPI-based nuclear staining. Then imaging of RNA seqFISH+ as well as sequential immunofluorescence for some targets was performed with the sequential hybridization and imaging routines described below. The samples were then disconnected from the fluidics system and proceeded to the DNA seqFISH+ preparation (see ‘Cell culture experiment’ and ‘Tissue slice experiment’). Next, the registered FOVs for RNA seqFISH+ were loaded and manually shifted to find the same cells in the original FOVs, followed by DNA seqFISH+ and sequential immunofluorescence imaging using the sequential hybridization and imaging routines described below.

The sequential hybridization and imaging routines were performed at room temperature on the automated confocal microscope. Briefly, for the sequential hybridization routine, the serial hybridization buffer, which consisted of a mixture of two or three unique readout probes (10-250 nM) with different fluorophores (Alexa Fluor 647, Cy3B or Alexa Fluor 488) and 10% EC buffer (10% ethylene carbonate (Sigma E26258), 10% dextran sulfate (Sigma D4911) and 4× SSC), was picked up from a 96-well plate and incubated in the flow cell for 20 minutes. The samples were then washed with 1 mL of a 4× SSCT buffer (4× SSC and 0.1% Triton-X), 330 μL of the 12.5% wash buffer, and 200 μL of 4× SSC, followed by a staining with about 200 μL of the DAPI solution consisting of 5 μg/mL DAPI (Sigma D8417) and 4× SSC for 30 seconds. The sample was then imaged with an anti-bleaching buffer consisting of 50 mM Tris-HCl pH 8.0 (Invitrogen 15568025), 4× SSC, 3 mM Trolox (Sigma 238813), 10% D-glucose (Sigma G7528), 100-fold diluted catalase (Sigma C3155), 1 mg/mL glucose oxidase (Sigma G2133). After the image acquisition detailed below, the sample was washed with 1 mL of the 55% wash buffer for 1 minute to strip off readout probes, followed by an additional incubation for 1 minute and rinsing with 4× SSC. Those serial hybridization, imaging, and signal extinguishing routines were repeated until the completion of all designated rounds.

The imaging conditions were determined based on the previous studies^7, 8^. Briefly, snapshots were acquired per fluorescent channel per field of view with 250 nm z-steps over 6 μm for cell culture experiments and 12 μm for tissue slice experiments. The pixel size for x and y is 103 nm. RNA seqFISH+ imaging was performed with 635 nm, 561 nm, and 488 nm fluorescent channels by omitting a 405 nm fluorescent channel to prevent a potential damage on the nuclei prior to DNA FISH^8^ except for a DAPI alignment hybridization round in the end. The readout probes for fiducial markers were also included in the first 2 fluorescent channels to allow image registration at the subpixel resolution. For the tissue slice experiments, polyA staining was performed in the 488 nm fluorescent channel. DNA seqFISH+ imaging was performed with 635 nm, 561 nm, 488 nm, and 405 nm fluorescent channels with DNA seqFISH+ targets in the first 3 fluorescent channels and DAPI staining in the 405 nm fluorescent channel. The readout probes for fiducial markers were also included in the first 3 fluorescent channels to allow image registration at the subpixel resolution. Imaging for sequential immunofluorescence was similarly performed with 635 nm, 561 nm, 488 nm, and 405 nm fluorescent channels, staining primary antibody targets in the first 2 fluorescent channels, fiducial markers in the third fluorescent channel, and DAPI in the last 405 nm fluorescent channel. At the beginning and end of the imaging rounds, fiducial marker images only with fiducial markers were obtained in the first 3 fluorescent channels for image registration. Furthermore, at the end of all imaging routines, images were manually checked and problematic imaging rounds such as off-focus and intensity saturation were repeated.

### Nuclear and cytoplasmic segmentation

The 3D nuclear and 2D cytoplasmic segmentation for individual cells were performed with a generalist, deep learning-based segmentation method, Cellpose^86^. First, for the nuclear segmentation, aligned and scaled images with Lamin B1, H3K27me3, H4K20me3, and DAPI in cell culture experiments or with BRG1, mH2A1, and DAPI in cerebellum tissue slice experiments were combined and used as 3D segmentation inputs for the Cellpose with specific parameters (flow_threshold=0.5, cellprob_threshold=0.5 in cell culture and flow_threshold=0.8, cellprob_threshold=0.5 in tissues). The original 3D nuclear segmentation labels were then eroded by 2 pixels if different labels were located at adjacent pixels to avoid potential misassignment. In addition, the labels whose centroids were located within 20 pixels from the edge of the images in x or y dimensions were filtered out. Second, for the cytoplasmic segmentation in cell culture experiments, aligned polyA images with a maximum intensity z-projection by ImageJ were used as 2D segmentation inputs for the Cellpose. The obtained cytoplasmic labels were then converted to ImageJ ROIs, eroded by 4 pixels, and manually corrected if necessary by using an ImageJ plugin, LabelsToROIs^87^. For cerebellum tissue slice experiments, we only computed nuclear labels and did not create cytoplasmic labels, similarly to our previous tissue slice experiments^8^. Finally, the obtained nuclear labels and cytoplasmic ROIs were loaded to MATLAB, and then were matched and renumbered by comparing the centroid location of nuclear labels with cytoplasmic ROIs. In this step, any matches without unique pairs were filtered out. The final nuclear and cytoplasmic labels for individual cells were stored as labeled images.

### seqFISH image processing

The seqFISH image processing steps were performed based on previous studies^7, 8^ with modifications. In brief, the preprocessing of the images were first performed by applying a flat field correction, a chromatic aberration correction considering full affine shifts in the X and Y dimensions (scaling, rotation, translation and shearing) but only translation in the Z dimension, and a background subtraction using the ImageJ rolling ball background subtraction algorithm with a radius of 3 pixels. Second, pixel locations for seqFISH spots were identified by using a Laplacian of Gaussians filter and a 3D local maxima finder with thresholding values obtained from semi-manual steps^7^. The identified seqFISH spot locations were further super-resolved at a sub-pixel resolution using a 3D radial center algorithm^88, 89^. Third, images from different hybridization rounds were aligned to the initial hybridization (hybridization 1) image in DNA seqFISH+ at a subpixel resolution by computing the translation of identified fiducial markers in each fluorescent channel. To align RNA and DNA images, the alignment to correct any rotation computed from DAPI staining images was further applied.

Using the aligned spots, seqFISH decoding was performed. The decoding of non-barcoded RNA seqFISH was performed based on the previous studies^7, 8^. In addition, the decoding of mRNA and intron seqFISH+ was performed similarly to those used for 1Mb-resolution DNA seqFISH+ experiments^7, 8^ with appropriate pseudocolor numbers and barcode keys with a voxel search radius of square root of 3 and one round of the error correction. We then used seed information generated during the barcode finding in seqFISH experiments for stringency^7, 8, 16, 17^. For mRNA seqFISH+ for cell culture experiments, we compared seed 3 and seed 4 results of mESCs, and filtered out top 150 barcodes, including on and off targets, that are differentially identified between those. Then the remaining seed 3 results were used for the downstream analysis. For intron seqFISH+ experiments, we used the results of at least 4 out of 5 seeds as before^16^. For the intron seqFISH+ experiments in cell culture, the distance to the nearest chromosome territory defined by the DNA seqFISH+ result was computed for each intron spot. Then intron spots within 500 nm from their own chromosome territories were considered to be transcription active sites (TASs), and other spots, which include introns outside TASs as well as false positives, were filtered out from the downstream analysis.

The decoding of two-layer DNA seqFISH+ was newly developed by leveraging the previous 25-kb resolution DNA seqFISH+ decoding approach^7, 8^. At each rounded pixel location where the spots were identified in the first 60 hybridization rounds, z-scored chromosome paint intensities for each cell and each hybridization round were computed for the next 36 hybridization rounds to provide 36 z-scored chromosome paint intensity values on each spot, corresponding to 9 pseudocolors for 4 barcoding rounds described in ‘Two-layer DNA seqFISH+ encoding strategy’. The barcodes for each spot were identified by matching the pseudocolors with the largest chromosome paint intensity z-score values in each barcoding round and compared with codewords identifying their genomic loci. To avoid false assignments, we dropped loci whose lowest chromosome paint intensity z-score in any barcoding round was above 0, or whose highest was below 0.5.

### Sequential immunofluorescence and DNA FISH image processing

To obtain subnuclear structures for individual cells, we used sequential immunofluorescence and repetitive DNA FISH images. Similarly to image processing for the spot detection described above, the images were corrected for a chromatic aberration shift and aligned to the initial hybridization image in DNA seqFISH+ by computing and propagating the translation of identified fiducial markers in an orthogonal fluorescent channel (488 nm) that was not used for sequential immunofluorescence or repetitive DNA FISH. In contrast to image processing for the spot detection, the background subtraction processing was not applied to the images. The mean intensities for each nuclear label were computed for all markers using the aligned images.

### Conversion of voxel information to physical size

After image processing steps above, we converted the voxel information of the images or decoded spots to physical size (0.103 μm for x and y and 0.250 μm for z) for the downstream analysis below.

### Separation of homologous chromosomes

In our previous works^7, 8^, we used DBSCAN to cluster genomic loci into their separate chromosome copies based on their spatial location. This worked well in most cases, but failed in cases where two copies of a chromosome were close to each other. To address these cases, we developed a package, DNASeqFISHChromosomeAssignment.jl, that separates loci into separate chromosome copies first with DBSCAN, then refines the clustering by taking each loci’s spatial and genomic coordinates into account. It checks whether each DBSCAN cluster has a proportion of unique loci below a threshold and whether the average spatial distance between subsequent loci is above a threshold. If so, it splits the cluster using an algorithm that we call Longest Disjoint Paths (LDP). This algorithm conceptualizes chromosomes as invisible strings on which we can image beads with DNA seqFISH. We can infer where the chromosomes are by connecting the dots between subsequent genomic loci to find a distinct path for every chromosome copy.

Finding the multiple longest disjoint paths is an integer programming problem similar to a maximum flow problem. To set up the problem, the LDP algorithm first constructs a directed graph where nodes represent loci. Loci within a user-specified spatial radius (different for different datasets) and different genomic coordinates are connected by directed edges pointing towards the locus of larger genomic coordinates. Edges are weighted according to the formula

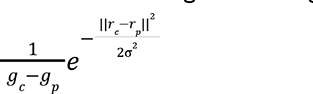

We then reduce the number edges in the graph by finding its transitive reduction, then removing any outgoing edge from a node that has lower weight than that node’s lowest weighted in the transitive reduction. To finish framing the problem, we add imaginary source and destination nodes to the graph that respectively have outgoing or incoming edges of no weight connecting them to every genomic locus node.

Our integer programming problem is tasked to find the maximally weighted edges that comprise one or more disjoint paths. To enforce that all edges are chosen are part of disjoint paths, we add the following constraints: the solution can choose either exactly one incoming edge and one outgoing edge, or no edges from each genomic locus node, and no more outgoing edges from the source than the maximum allowed number of chromosomes that can be chosen.

### 2D and 3D visualization of DNA seqFISH+ data

The 2D visualization of DNA seqFISH+ spots was performed as described before^39^. Because raw DNA seqFISH+ images are barcoded and the identities of DNA spots are indistinguishable without decoding, we reconstructed images after decoding from rounded voxel location of the decoded DNA seqFISH+ spots by applying a multidimensional Gaussian filter (sigma = 1) with scipy.ndimage.gaussian_filter package in python and overlaid with aligned raw immunofluorescence images by using ImageJ. Those composite images are displayed in 2D either as a single z-section or a maximum z-projection of multiple z-sections as specified in each figure caption.

The 3D visualization of DNA seqFISH+ or intron seqFISH+ data was performed by using python API Mayavi2^90^. Physical coordinates of all detected DNA spots or decoded intron spots were stored as x, y and z numpy arrays and then visualized as spheres. DNA spheres were colored differently to represent different chromosomes or gene families, or displayed as transparent small spheres when they were shown as background. Intron spheres were colored by chromosome identities of each gene. Genomically adjacent DNA spots were connected using tubes to show the organization of a single chromosome.

H4K20me3 immunofluorescence and MajSat DNAFISH signals were visualized by displaying a surface around x, y and z coordinates with intensity z-score values above 2.

### Sequencing feature analysis in the mouse genome

To compute GC content per 25 kb genomic bins, we created a bed file for the 25 kb bins with unmasked mouse mm10 reference genome and performed the computation within each bin using ‘bedtools nuc’^91^. To compute the gene density and gene length features per 25 kb genomic bins, we downloaded GRCm38/mm10 refGene database, retrieved the longest gene annotation for each gene, and mapped back the gene name and corresponding gene length to each 25 kb genomic bin. We then further computed the GC content, gene density per 100 kb and 200 kb bins. Meidan gene length per 100 kb and 200 kb bins were defined by the median gene lengths of genes falling in those bins.

### Sequencing-based data analysis

A/B compartment score file from the Hi-C matrix was downloaded from the original study at 100 kb resolution for mESCs^20^ or kindly provided by Sandra Peiró at 25 kb resolution for NMuMG cells^67^.

Compartment scores of NMuMG cells were further binned using a mean score of 25 kb bins in each 100 kb bin.

Processed bulk RNA-seq data for cell culture were obtained from NCBI GEO (accession GSE98674 for mESCs^66^ and accession GSE96033 for NMuMG cells^67^). For the GRO-seq analysis in mESCs^68^, single-end FASTQ files were downloaded from NCBI GEO (accession GSE48895) and first trimmed to keep the first 32 bases using the cutadapt package (version 1.18)^92^. Trimmed reads were aligned to the mm10 genome using bowtie2^93^. Uniquely mapped reads were filtered and sorted before duplicate reads were removed using samtools^94^. Strand-specific RPKM bigwig signal files were generated using bamCoverage with filterRNAstrand option employed and step size of 10. Forward and reverse strand aligned gene body bed files (generated from GENCODE vM25 annotation) were used to compute GRO-seq signal intensity within gene body regions using computeMatrix command of deeptools. GRO-seq intensity within ±500bp around gene transcription start sites was calculated using a similar approach.

The bigwig CUT&Tag datasets for H3K27ac, H3K27me3, H3K4me1, and H3K4me3 in mESCs were obtained from NCBI GEO (accession GSE166210)^70^. Average bigwig signals across 25 kb genomic bins were calculated using ENCODE bigWigAverageOverBed software. Bin ID and calculated score were extracted. Average bigwig signals were further average binned to 100 kb genomic bins for validation.

The processed DamID dataset for Lamin B1 in mESCs was obtained from NCBI GEO (accession GSE181693)^69^ and binned with 100 kb resolution. The different lamina-associated domain categories (cLADs, fLADs, and ciLADs) from Lamin B1 DamID datasets were obtained from NCBI GEO (accession GSE17051)^71, 72^ and mm9 genomic coordinates of the obtained files were converted to mm10 using the UCSC Genome Browser program liftover.

### Differential gene expression analysis

For cell culture study, single cell intron count matrices and metadata from mESC and NMuMG intron seqFISH+ experiments were loaded to Seurat (version 3.2.0)^95^. Intron counts of two biological replicates of mESC sample and one biological replicate of NMuMG sample were aggregated to generate normalized ensemble count data for mESC and NMuMG respectively. Intron differential expression analysis between mESC and NMuMG were performed using DESeq2^96^ (version 1.26.0). Genes with the adjusted p-value <0.01 and absolute value of log2 fold change > 2 were selected as significantly differential expressed genes.

The adult mouse cerebellum single-nucleus RNA-seq (snRNA-seq) data^26^ was obtained from NCBI GEO (accession GSE165371). Single-cell cell-gene count matrices and sample meta tables were loaded to Seurat (version 3.2.0) for cell-type specific analysis. Gene counts of each cell cluster were aggregated for every replicate sample as ensemble count data for differential expression analysis using DESeq2.

Differentially expressed genes were selected using the same criteria described above. Transcriptional activities across the mm10 reference genome were computed by summing averaged DEseq2 normalized counts of identified differentially expressed genes located within each 200 kb genomic bins within cell types. The resultant average gene counts at 200 kb bins were used to calculate log2-fold expression change between pairs of cell types. The same analysis was performed by using intron seqFISH+ measurements with cell type information defined by mRNA seqFISH clusters.

For the cerebellum study, to compare the relationships between cell-type specific gene expression and immunofluorescence marker enrichment, differentially expressed 200 kb bins between pairs of cell types obtained above were selected, and the ensemble-averaged immunofluorescence marker differences between the same cell type pair were further calculated, similarly to our previous study^8^. Box plots were then used to show the immunofluorescence marker differences between those differentially expressed 200 kb bins computed from intron seqFISH+ measurements.

### Imaging-based transcriptomic data analysis

The similarity of ensemble-averaged mature or nascent transcriptome profiles by mRNA seqFISH+ or intron seqFISH+ in cell culture was compared to those by bulk RNA-seq in mESCs^66^ and NMuMG cells^67^ or GRO-seq in mESCs^68^ by using Spearman correlation, confirming high consistency of the datasets.

The preprocessing, clustering, and visualization of the imaging-based transcriptomic data (mRNA and intron seqFISH+) for the adult mouse brain cerebellum were performed with a Scanpy toolkit^97^ in Python. For mRNA seqFISH analysis, we chose genes based on the max copy number of at least 20 in any cells, which yielded 49 genes. The gene count matrix with those genes was then used for the further clustering analysis. We then applied total count normalization and log(counts + 1) to transform the matrix, and performed the dimensional reduction with principal component analysis (PCA), followed by batch correction with Scanorama^98^. We then constructed a nearest-neighbor graph with k = 40 neighbors using the top 30 principal components, followed by clustering of cells with Leiden clustering^99^ and embedding using uniform manifold approximation and projection (UMAP)^100^. Similarly, cells were clustered with Scanpy^97^ using intron seqFISH+ datasets. Briefly, we applied total count normalization, log(counts + 1), and scaling to transform the matrix, and performed the dimensional reduction with PCA and batch correction with Scanorama^98^. We then performed construction of a nearest-neighbor graph with k = 25 neighbors using the top 30 principal components, Leiden clustering^99^, and UMAP embedding^100^. During these steps we filtered out cells based on the nuclear volume. Specifically, we filtered out cells with nuclear volume less than 100 μm^3^ as a potential mis-segmentation. In addition, after Leiden clustering, we filtered out cells with nuclear volume outside the interquartile range in each cluster to minimize the doublets. In the end, we obtained *n* = 4,015 cells, consisting of *n* = 1,504, 832, 518, 357, 263, 164, 113, 88, 76, 56, 29, 15 cells from Leiden cluster 0 to 11, in two biological replicates of adult mouse brain cerebellum datasets (Supplementary Table 3).

The clustering results for individual cell types of the mouse brain cerebellum were compared to the normalized pseudo-bulk snRNA-seq expression profiles of each cell type in the adult mouse cerebellum, computed in the original study^26^. The gene expression profiles of 49 genes after the filtering described above were used for the comparison. The degree of similarity for each cell type between imaging and sequencing datasets was evaluated by using the Pearson correlation and cell-type identity was annotated to each mRNA seqFISH cluster. Those clusters were annotated as Granule (cluster 0, 1), Bergmann glia (cluster 2), MLI1 (cluster 4), Purkinje cells (cluster 6, 11), MLI2/PLI (cluster 7), Endothelial (cluster 8), Astrocyte (cluster 9), and OPC/ODC (cluster 10). Clusters 3 and 5 were filtered out due to the mixture of cell types. Similarly, nascent transcriptional profiles of the four major cell types (Granule, Bergmann glia, MLI1, and Purkinje cells) in the adult mouse brain cerebellum by intron seqFISH+ were compared to RNA expression profiles of each cell type obtained from snRNA-seq datasets^26^, using a subset of genes with detected average copy number of more than 0.1 per cell per each cell type by intron seqFISH+.

### Single-cell global chromatin state analysis

Cells were clustered by averaged intensity profiles of individual immunofluorescence markers in each nucleus as performed before^8^ by using a Scanpy toolkit^97^ in Python. Briefly, we used the mean intensity profiles of each immunofluorescence marker (*n* = 27 markers) within each nuclear label. Similarly to the intron seqFISH+ analysis, we then applied total intensity normalization, log(intensity + 1), and scaling to transform the matrix, and then performed the dimensional reduction with PCA and batch correction with Scanorama^98^. Then we performed construction of a nearest-neighbor graph with k = 40 neighbors using the top 30 principal components, Leiden clustering^99^, and UMAP embedding^100^. The similarity of obtained clusters was compared to those from mRNA seqFISH by computing the overlapped fraction between a given pair of clusters.

### Estimation of detection efficiency and false positive rates for two-layer DNA seqFISH+

The estimation of detection efficiency by two-layer DNA seqFISH+ for cycling mESCs as well as post-mitotic diploid cells in the female mouse brain was performed similarly to our previous studies^7, 8, 101^. Briefly, by considering the cell cycle distribution, we estimated detection efficiency as 21.1 ± 6.8% (median ± s.d.) from 63,466 ± 20,525 (median ± s.d.) DNA spots per cell for 100,049 loci in single male mESCs (n = 1,076 cells), while 5.0 ± 3.5% (median ± s.d.) from 9,912.0 ± 6,932.0 (median ± s.d.) counts per cell for 100,049 loci in post-mitotic Purkinje cells (n = 113 cells) in the female mouse brain cerebellum. We note that the lower detection efficiency of the brain samples could be caused by incomplete coverage of entire nuclei in z direction for some cells, optical clouding of spots due to smaller nuclear sizes in tissues, as well as reduced hybridization efficiency due to the tissue thickness and different fixation conditions.

The estimation of false positive rates from two-layer DNA seqFISH+ was adapted from the previous DNA seqFISH+ scheme^7, 8^. To compute the false positive rates, both on-target barcodes (n = 100,049 barcodes) and blank barcodes, consisting of all the remaining error-correctable barcodes (n = 31,171 barcodes) in the codebook, were run simultaneously. The false positive rates were then computed by median on-target barcode counts per barcode per cell divided by median total barcode counts per barcode per cell, which provided false rates of 1.2% in mouse ESCs (n = 1,076 cells) and 4.3% in the adult mouse brain cerebellum (n = 4,015 cells).

### Pairwise spatial distance analysis for DNA loci

We calculated mean pairwise distances between genomic loci similarly to our previous studies^7, 8^ with modifications. When computing pairwise distances for one chromosome, we only considered distances between pairs of loci that we found to be on the same homologous chromosome. To calculate mean distances, we count co-detections and sum pairwise distances for each loci pair in separated chromosomes in each cell. After evaluating for all cells, we divide the sums of pairwise distances by the number of co-detections. The pairwise spatial distance maps without binning (25-kb) or with 200-kb or 1.5-Mb (paint barcode) binning were used as specified in each figure.

The 1-Mb and 25-kb DNA seqFISH+ datasets (*n* = 2,460, 1,200 loci, respectively) from two biological replicates of mESCs^7^ were obtained from the 4D Nucleome data portal (https://data.4dnucleome.org/) under accession number 4DNESL2AY9CM. The median spatial distances were then compared to those from this study by using commonly profiled genomic loci. The two-layer DNA seqFISH+ spatial distance maps typically contain boundaries between loci encoded in different chromosome blocks in different fluorescent channels, possibly due to the the grouping of the DNA loci into blocks and the differences in the co-detection efficiency between loci across different blocks. We note that these boundaries do not affect all the analysis performed in the paper, such as compartment analysis, spatial chromatin profiling, and spatial distance analysis (>= 1 Mb resolution), which were further validated by orthogonal datasets in mESCs.

Characterization of inter-chromosomal interaction was conducted at the 1.5-Mb (paint barcode) resolution distance matrix. Spatial distances between top 5% genomic loci associated with each subnuclear marker or all loci annotated with specific gene families were selected, and distances from pairs of intra-chromosomal loci were filtered out. Random 5% of the genomic loci were selected by bootstrap 1,000 times and inter-chromosomal spatial distances from random selected genomic loci were extracted to compare with marker specific spatial distances.

### Radial organization analysis for DNA loci within the nucleus

The radial organization of DNA loci within the nucleus was evaluated by computing the convex hull surface of the DNA loci similarly to previous studies^5, 6, 102^. To compute the radial positioning of DNA loci within the nucleus, we constructed a 3D convex hull for each nucleus using the DNA seqFISH+ spots per cell using the SciPy spatial library in Python. At this step, cells with less than 100 DNA seqFISH+ spots were filtered out. We then calculated the spatial distance of individual spots from the nuclear periphery by calculating the distance of intersection from the convex hull surface to the centroid of the nucleus through the individual spots. The radial scores of genomic loci were similarly computed by scaling the spatial distance from the nuclear center to nuclear periphery as 0 to 1. The computed median distance profiles from the nuclear periphery in each cell type were compared by Pearson or Spearman correlation with Lamin B1 chromatin profiles of the corresponding cell type, which are independently measured and typically enriched at the nuclear periphery, with 100-kb binning for the cell culture datasets and with 200-kb binning for the adult mouse brain cerebellum datasets.

The similarity of radial positioning of genomic loci among cell types in the adult mouse brain cerebellum was computed by Pearson correlation of median radial scores across cell types with 200-kb binned loci. The top 5% variable loci in radial positioning between neurons and glial cells were computed by using the radial profiles of neuronal cells (Purkinje cells, MLI1, MLI2/PLI) and glial cells (Bergmann glia, Astrocytes) as input. Out of 629 variable loci, we identified 409 loci moved interior of the nucleus in neurons versus 220 loci in glial cells. We then compared the degree of overlap between those variable loci and Purkinje H4K20me3-associated loci or mCH desert loci^48^.

### Imaging-based chromatin profiling analysis

We used z-score normalization of each chromatin marker (antibodies and ncRNAs) in each 3D nuclear label to compute voxel resolution chromatin profiling, similarly to those previously described ^7, 8^. Briefly, the intensity values for each marker obtained from aligned sequential immunofluorescence or RNA FISH or DNA FISH images were normalized by computing intensity z-scores at individual voxels per each nuclear label and then the z-scored intensity profiles for each marker were computed at rounded voxel locations where the final decoded DNA spots were identified by DNA seqFISH+. This approach allows us to investigate matrices consisting of DNA loci by subnuclear markers in individual cells. Specifically, each single cell was presented as DNA loci to subnuclear marker matrix, where each detected 25-kb DNA locus is represented as a vector with features including the normalized z-scores of subnuclear marker intensities, as well as radial positioning inside the nucleus.

Then DNA loci and their associated features were binned into 100-kb, 200-kb, or 1.5-Mb (paint barcode) resolutions. Single cells were then grouped according to cell type to calculate the ensemble-averaged level of subnuclear marker scores per each cell type. For cell culture study, median scores of each feature associated with DNA bins from mESCs or NMuMG cells were calculated to get the ensemble-averaged marker intensity z-score at 25-kb, 100-kb, 200-kb or 1.5-Mb resolution. For the cerebellum data, single cells assigned to different cell types were grouped and median scores of each feature were calculated per each cell type at 200-kb and 1.5-Mb resolution. For both cell culture and cerebellum studies, the rare case of DNA bins which were filtered out during radical score calculation were also filtered out in this analysis.

To validate the imaging-based chromatin profiles in this study, we compared those with 1-Mb resolution imaging-based chromatin profiles by DNA seqFISH+ obtained from the original study^7^ and other sequencing-based datasets such as CUT&Tag and DamID^69, 70^. The comparison of chromatin profiles between DNA seqFISH+ (*n* = 2,460 loci) and two-layer DNA seqFISH+ was performed at commonly profiled 25-kb genomic loci that are on average ∼1 Mb apart, while those between two-layer DNA seqFISH+ and sequencing datasets were with 100 kb binning. The degree of similarity between datasets was evaluated by Pearson or Spearman correlation. In addition, to further validate the genomic loci to the nuclear lamina association, we compared the enrichments of imaging-based Lamin B1 profiles from mESCs across previously identified lamina-associated domains^71, 72^. See ‘Sequencing-based data analysis’ for the details of sequencing-based data source and processing.

The similarities of genome-wide chromatin profiles between cell types were evaluated by computing the Pearson correlation coefficient of each marker between given pairs of cell types with 200 kb binning. In addition, the similarities of chromatin profiles between pairs of markers in each cell type were evaluated by comparing the overlap of top 5% genomic loci associated with each marker in each cell type.

To compare differential association between two markers in each cell type, the top 5% differentially associated bins were selected. The ensemble-averaged z-score intensity of two markers were plotted along the x- and y- axis, respectively. We then moved a slope = 1 line along the x or y axis until 5% of the bins above or lower this line were identified. Similarly, SF3A66 z-score change and RNAPIISer5-P z-score change between two cell types were plotted along the x- and y-axis, respectively. A slope = 4 line was moved along the SF3A66 axis until 5% of the bins were on the left or right side of this line to define the top 5% SF3A66 differentially associated bins. A slope = 0 line was moved along the

RNAPIISer5-P axis until 5% of the bins were above or below the line to define the 5% RNAPIISer5-P differentially associated bins.

We also computed compartment scores^103^ by using selected immunofluorescence markers including H3K27ac, H3K27me3, H3K9me3, and Lamin B1 in mESCs and NMuMG cells. For each chromosome, correlation matrices of mESC and NMuMG 100 kb ensemble-averaged immunofluorescence z-scores for the above 4 markers were computed. The first eigenvectors of the above correlation matrices were calculated as immunofluorescence A/B compartment scores. The sign of the compartment scores was checked for each chromosome and multiplied by -1 to flip the signs if necessary to correct the direction of the eigenvector of the entire chromosome. The similarity of Hi-C and imaging datasets were then compared by computing Spearman correlation at 100-kb resolution.

### Peak definition of chromatin profiles

To characterize genome fragments associated with various subcompartments, peak calling was performed on ensemble-averaged 200-kb binned z-score tracks for both cell culture and cerebellum studies. One dimension ensemble-averaged z-scores of each subnuclear marker were mean-centered within chromosomes first, then scipy.signal.find_peaks function was performed with manually decided “prominence” and “height threshold” parameters to call out the peaks. Overlapping peaks were merged to one broader peak. For the nuclear periphery distance, negative mean centered distance was used due to its negative correlation with lamina-associated domains. Genomic bins characterized as Purkinje H4K20me3-associated peaks (see ‘Characterization of H4K20me3-associated genomic loci’) were compared to the peak definition in other cell types for H4K20me3 and nuclear periphery. Then the transition of H4K20me3 bins into other type of repressive bins in other cell types were visualized using Sankey plot using pySankey package.

To classify active subnuclear compartments, RNAPIISer5-P peaks spanning more than 2-Mb were defined to be broad peaks and rest peaks were defined to be sharp peaks. SF3A66 peaks overlapped with RNAPIISer5-P peaks were assigned to be speckle peaks. Similarities of speckle peaks, broad and sharp RNAPIISer5-P peaks across cerebellum cell types were calculated by defined overlapping score: the sum of the fractions of the number of overlapping peaks in all peaks in both cell types and then divided by 2. To compare to the active subnuclear compartments, we computed repressive peaks using peaks at least in one of the following repressive compartments (nuclear periphery, H4K20me3, or H3K27me3).

### Characterization of cell-type-specific long genes

To characterize long gene specific association with nuclear compartment, genes with > 200 kb were defined as long genes and DNA bins containing long genes were selected for further analysis.

Pseudo-bulk normalized mRNA counts of different cell types in the adult mouse brain cerebellum provided by snRNA-seq study^26^ were used to represent mRNA expression level. For each long gene, the maximum RNAPIISer5-P normalized z-scores among the gene spanned 200kb DNA bins were calculated as long gene RNAPIISer5-P level. Pearson correlation between normalized mRNA counts and RNAPIISer5-P z-scores of each long gene across cell types was calculated. Among 867 defined long genes, 132 and 280 genes, which have a correlation higher than 0.9 and 0.8 respectively, were identified. To further evaluate other subnuclear marker profiles in long gene regions, jaccard index between peaks of other subnuclear markers and RNAPIISer5-P peaks in long gene DNA bins were calculated using scipy.spatial.distance. Specific immunofluorescence markers (SOX2, HDAC1, ATRX, Fibrillarin, RNAPIISer2-P) were filtered out from this analysis due to the cell-type specific staining or low signal-to-noise ratio.

To compare the relationship between RNAPIISer5-P enrichment and open chromatin region at the long gene loci, the processed file of genome length for the open chromatin regions by ATAC-seq in Purkinje cells was obtained from the original study^77^. Then the Purkinje RNAPIISer5-P enrichment at the cell-type-specific RNAPIISer5-P long gene loci (Pearson’s *r* > 0.8) with or without a highly open conformation containing >20,000-bp peaks^77^ (n = 17, 263 genes, respectively) were compared by two-sided Wilcoxon’s signed rank-sum test. We note that the cell-type-specific RNAPIISer5-P long genes contain genes expressed in Purkinje cells as well as other cell types as described above.

To identify Purkinje-specific H3K27me3-associated long genes, the cell-type-specific H3K27me3-associated loci were first identified by comparing H3K27me3 profiles between pairs of cell types in the adult mouse brain cerebellum (see ‘Imaging-based chromatin profiling analysis’). Then Purkinje-specific H3K27me3-associated long genes (*n* = 116 genes) were obtained by finding long genes (>200 kb) annotated in the Purkinje-specific H3K27me3-associated loci relative to MLI1, MLI2/PLI, or Bergmann glia cells. The developmental gene expression profiles of those long genes were further examined by using a differential gene expression analysis with a provided processed data^46^, identifying 36 out of 46 developmental differentially expressed genes, which is a subset of Purkinje-specific H3K27me3-associated long genes, are down-regulated from P0 to adult Purkinje cells.

### Characterization of H4K20me3-associated genomic loci

To examine H4K20me3-associated genomic locus, 200-kb DNA bins identified as H4K20me3-associated peaks that are not assigned as MajSat peaks in the analysis under ‘Peak definition of chromatin profiles’ were further processed. H4K20me3-associated DNA bins with a normalized z-score >= 1 in Purkinje cells were defined to be H4K20me3-associated bins, others are defined to be weak H4K20me3-associated bins. To compare the other repressive marker enrichments at the H4K20me3-associated loci in Purkinje cells, quantile-normalized immunofluorescence marker z-scores were compared.

To characterize gene families and functions associated with H4K20me3 subcompartment, gene prefixes were extracted by cropping gene name symbols before any digit or symbol character. Recurrent gene prefixes were considered to be gene family, and then enrichment of gene families in

H4K20me3-associated locus were computed at 200kb resolution. Number of H4K20me3-associated bins in Purkinje cells that overlap with DNA bins containing SSDR genes^50, 51^ or DNA bins previously identified as mCH desert^48^ were further counted by converting mm9 genomic coordinates provided by the original studies to mm10 using the UCSC Genome Browser program liftover.

Individual H4K20me3 profiles in Purkinje cells were further investigated to characterize the chromatin organization of individual alleles at strong and weak H4K20me3-associated DNA bins.

H4K20me3-associated peaks enriched with *Vmn* and *Olfr* gene families were selected to represent strong and weak H4K20me3-associate genomic locus respectively. The genomic region spanning up to 10 Mb upstream or downstream of the beginning or end *Vmn* or *Olfr* H4K20me3-associated peaks were selected to separate homologous chromosomes. H4K20me3 and Lamin B1 immunofluorescence intensity z-scores in those regions from all alleles detected in Purkinje cells were collected, then visualized as heatmaps.

### Gene ontology analysis

Gene ontology (GO) analysis was performed using ClusterProfiler (3.18.1)^104^ under R (4.0.3) platform. Specific pairs of inputs below were formed into a gene list with corresponding annotations, and comparison between different annotations were conducted using “compareCluster” function using default settings, then gene ontology terms were further combined using “simplify” function at 0.7 cutoff. Visualizations of pathway enrichment were plotted by calling the “dotplot” function built in ClusterProfiler.

To compare the pathway enrichment between mESC and NMuMG cells, GO analysis was performed on the differentially expressed genes identified by intron seqFISH+. To compare the speckle and RNAPII associated gene pathways, genes overlapping with top 5% of SF3A66 or RNAPIISer5-P genomic loci in each major cell type in the adult mouse cerebellum were selected to compare the pathway enrichment. To obtain pathway enrichment associated with H3K27me3, genes overlapping with top 5% genomic loci associated with H3K27me3 or Lamin B1 in each major cell type in the adult mouse cerebellum were used for GO comparison. Similarly, genes located in Purkinje H4K20me3 or H4K20me3 weak genomic loci were extracted and compared for the GO enrichment. 200 kb binned genomic loci were used in this analysis for the tissue experiments.

### Statistics and reproducibility

Cells shown in Fig. 1 and Extended Data Figs. 2-4 are representatives of 1,076 cells from two biological replicates of mESCs or 384 cells from one biological replicate of NMuMG cells. Cells shown in Figs. 2-6 and Extended Data Figs. 5-10 are representative of 4,015 cells from two biological replicates of the adult mouse brain cerebellum.

---

**Extended Data Fig. 1.**
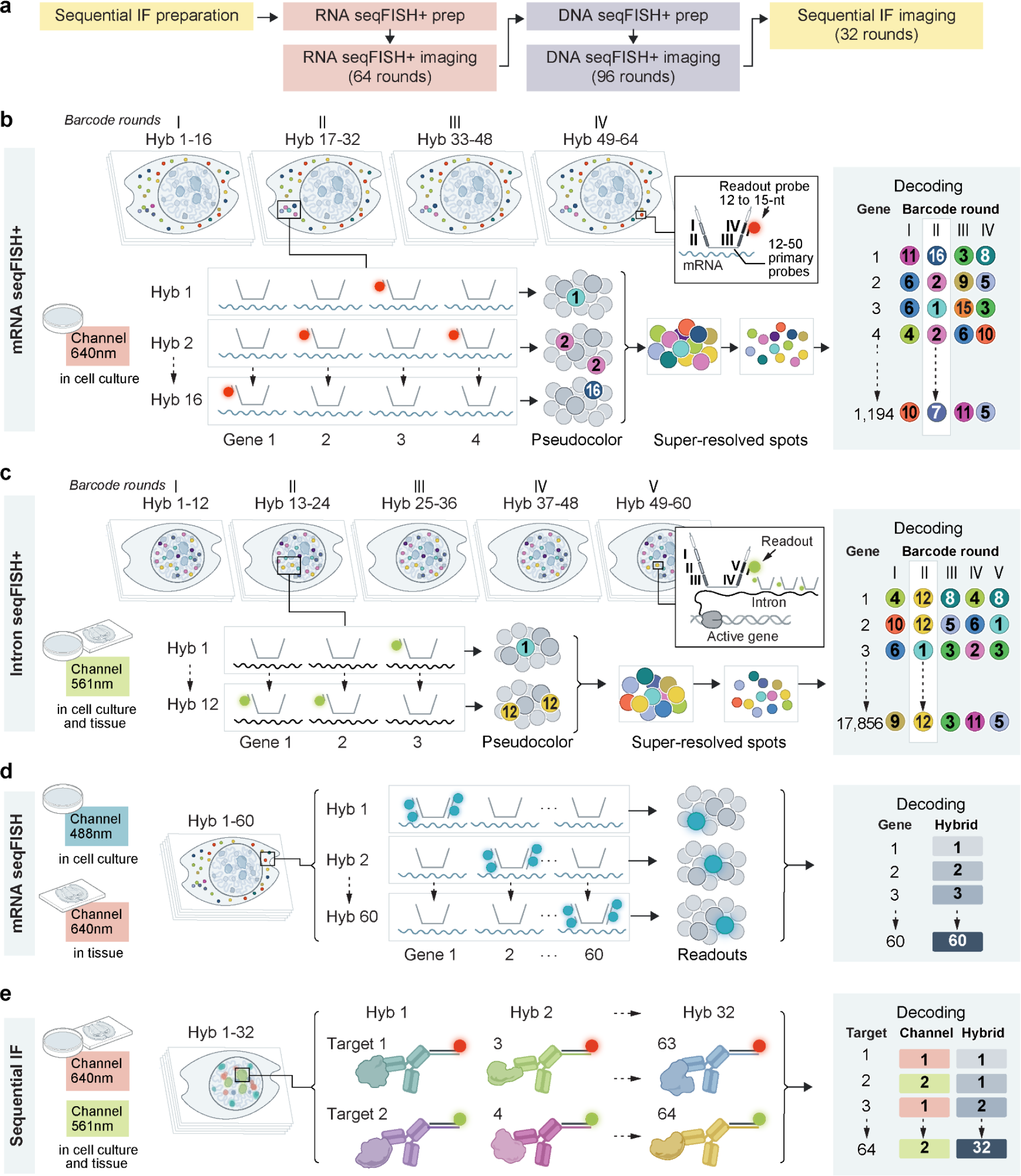
**Detailed schematic of single-cell spatial multi-omics. a**, Flow chart of the sample preparation and imaging in single-cell spatial multi-omics. **b**, Schematic of mRNA seqFISH+ with 4 barcoding rounds with 16-pseudocolors including one round of error correction using one fluorescent channel (640 nm). This coding scheme can resolve up to 4,096 barcodes, while a subset of 1,194 barcodes were used to resolve mRNA species in cell culture experiments. **c**, Schematic of intron seqFISH+ with 5 barcoding rounds with 12-pseudocolors including one round of error correction using one fluorescent channel (561 nm). This coding scheme can resolve up to 20,736 barcodes, while a subset of 17,856 barcodes were used to resolve intronic RNA species in cell culture and tissue experiments. **d**, Schematic of non-barcoded mRNA seqFISH using one fluorescent channel (488 nm in cell culture experiments; 640 nm in tissue experiments). Unlike the exponential barcoding scheme in **b** and **c** whose coding capacity increases exponentially to the number of barcoding rounds, the number of RNA species that can be distinguished increases linearly to the number of hybridization rounds. **e**, Schematic of sequential immunofluorescence using two fluorescent channels (640 nm and 561 nm) in cell culture and tissue experiments. Similar to **d**, the number of antibody species that can be multiplexed scales linearly to the number of hybridization rounds.

**Extended Data Fig. 2.**
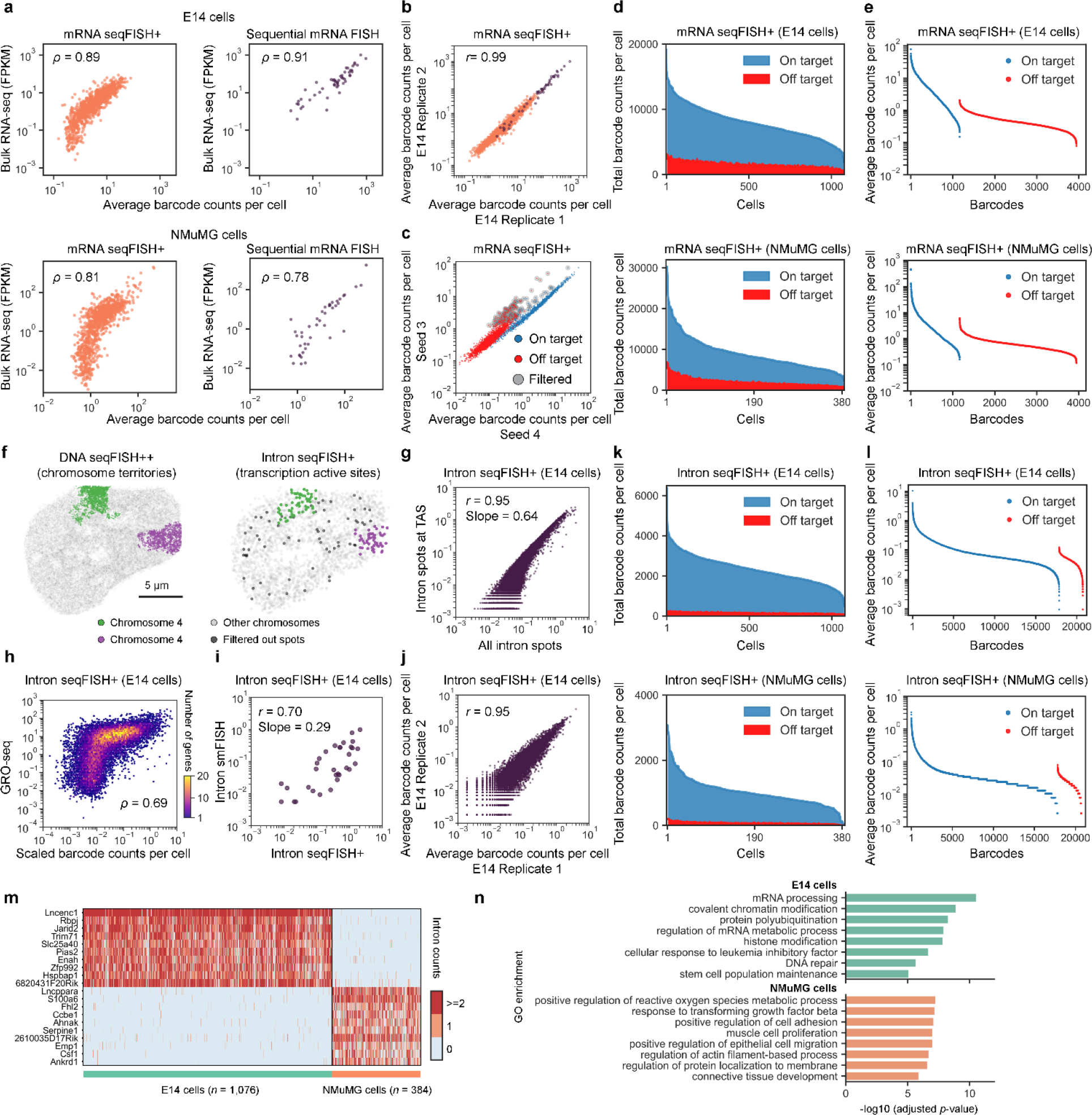
**Validation of mRNA and intron seqFISH+ in cell culture experiments. a**, Spearman correlation between average mRNA counts by mRNA seqFISH+ (left) or non-barcoded mRNA seqFISH (right) and bulk RNA-seq in mESCs^66^ (top) and NMuMG cells^67^ (bottom). **b**, Pearson correlation of average mRNA counts profiled by mRNA seqFISH+ (orange) and non-barcoded mRNA seqFISH (purple) between two biological replicates from mESCs. *n* = 1,194 and 53 genes for mRNA seqFISH+ and non-barcoded mRNA seqFISH profiling, respectively, in **a**, **b**. **c**, Visual representation of on- and off-target barcode counts and filtered barcodes by comparing mRNA seqFISH+ results from mESCs between seeds 3 and 4 decoding stringency (Methods). Those filtered barcodes (*n* = 150 barcodes, including both on- and off-target barcodes) were excluded from the downstream analysis. **d**, Total on- and off-target barcodes detected per cell by mRNA seqFISH+ in mouse ES cells (top) and NMuMG cells (bottom). **e**, Average counts per each on- and off-target barcode per cell by mRNA seqFISH+ in mouse ES cells (top) and NMuMG cells (bottom). *n* = 1,163, 2,783, and 150 for on-target, off-target, and filtered barcodes, respectively in **c**-**e**. **f**, Representative visualization of homologous chromosomes (chromosome 4) by DNA seqFISH+ and transcription active sites by intron seqFISH+ near chromosome territories in the nucleus of mESCs. Intron spots detected within 500 nm from chromosome territories of their own chromosome captured by DNA seqFISH+ were considered to be transcription active sites, and other spots were filtered out from the downstream analysis. **g**, Pearson correlation between average intron counts per cell computed from all intron spots and transcription active sites defined by the criteria in **f**. **h**, Spearman correlation between scaled intron counts by intron seqFISH+ and bulk GRO-seq^68^, measuring the amount of transcriptionally active RNA polymerase II (RNAPII), in mESCs. **i**, Pearson correlation of average intron counts at the transcription active sites by intron seqFISH+ and non-barcoded intron seqFISH for *n* = 33 genes^16^ in mESCs. The slope of 0.29 indicates a relative detection efficiency of 29%. **j**, Pearson correlation of average barcode counts per cell by intron seqFISH+ between two biological replicates from mouse ES cells. *n* = 17,856 genes for intron seqFISH+ in **g**, **h**, **j**. **k**, Total barcode counts per cell by intron seqFISH+ in mESCs (top) and NMuMG cells (bottom). **l**, Average counts per each on- and off-target barcode per cell by intron seqFISH+ in mESCs (top) and NMuMG cells (bottom). *n* = 17,856 and 2,880 for on- and off-target barcodes, respectively in **k** and **l**. **m**, Heatmap of gene expression profiles across single cells for top differentially expressed genes between mESCs and NMuMG cells by intron seqFISH+ (*n* = 10 genes for each cell line). **n**, The GO terms (top five plus manually selected three) identified from intron seqFISH+ profiles between mESCs and NMuMG cells represent their corresponding cell type identities. *n* = 1,076 cells from two biological replicates of mESCs and *n* = 384 cells from one biological replicate of NMuMG cells in **a**-**n**.

**Extended Data Fig. 3.**
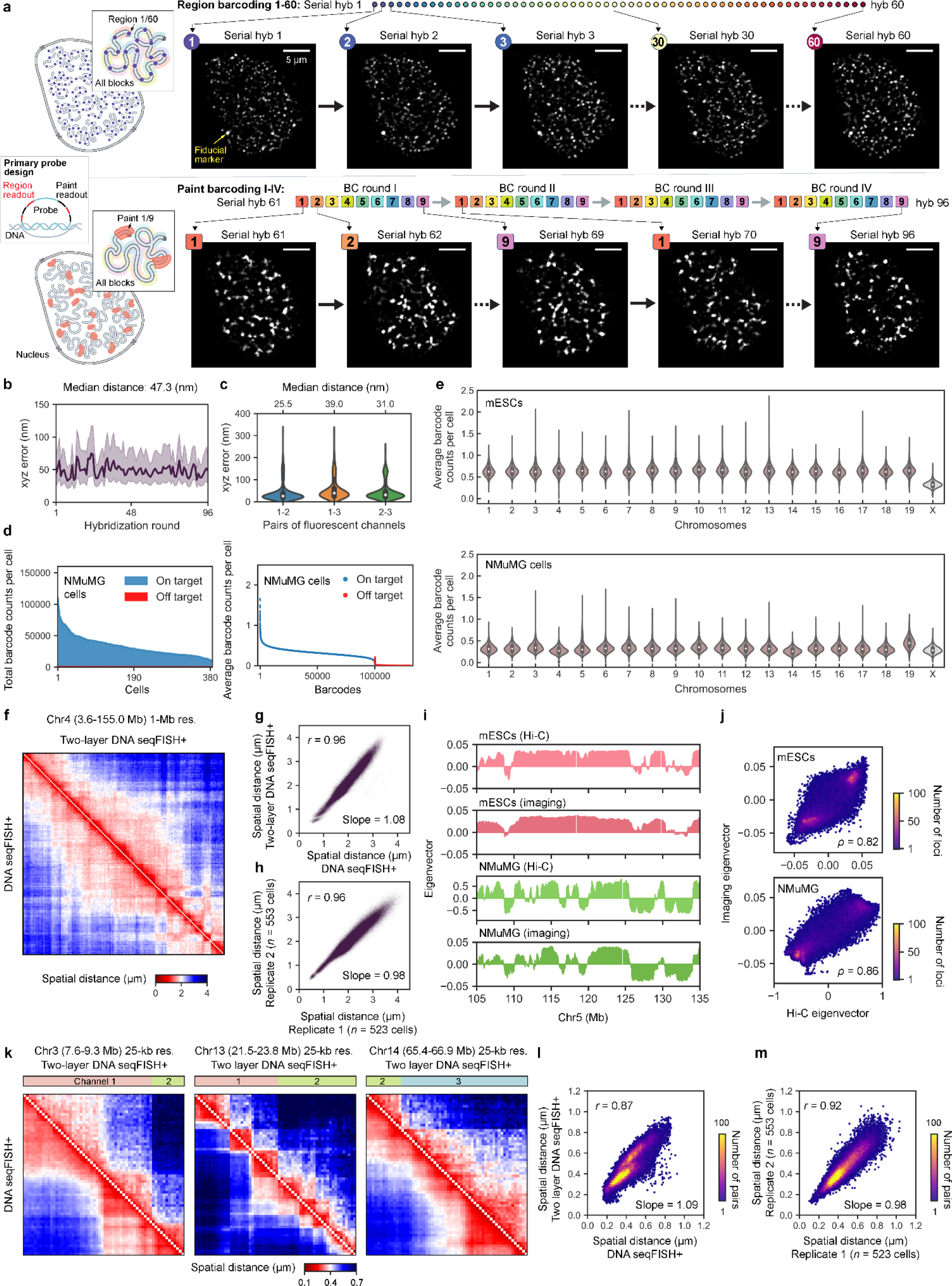
**Additional schematic and validation of DNA seqFISH+ in cell culture experiments. a**, Implementation of DNA seqFISH+ (left), by leveraging the combinations of region barcoding (top) and chromosome block barcoding (bottom). To implement this, each primary probe (left, middle) contained three identical 15-nt readout binding sites (black) for one of the 60 regions in hybridizations 1-60 as well as three 15-nt readout binding sites (red) for three out of four chromosome paint barcoding rounds (hybridizations 61-96) in each fluorescent channel. Representative images for the nucleus from mESCs across 96 rounds of serial hybridization and imaging of two-layer DNA seqFISH+ (right) for region barcoding (top) and chromosome block barcoding (bottom). A fiducial marker targeting a locally repetitive 3632454L22Rik locus by a single DNA FISH probe^7^ appears in all rounds of imaging. Background subtracted images used for an analysis (Methods) are shown from a single z-section for visual clarity. **b**, Quantification of the median fiducial marker localization relative to the reference image across 96 rounds of two-layer DNA seqFISH+ imaging, representing a high image alignment accuracy with median error of 47.3 nm in 3D across imaging rounds. Shaded areas represent the interquartile range. *n* = 1,332-2,402 matched fiducial markers in each hybridization round from in three fluorescent channels. **c**, Quantification of the median reference fiducial marker localization between pairs of fluorescence channels after the chromatic shift correction in 3D, suggesting a minimum chromatic effects across fluorescent channels after the correction. *n* = 2,301 matched spots. **d**, Total on- and off-target barcodes detected per cell by two-layer DNA seqFISH+ in NMuMG cells (left). Average counts per each on- and off-target barcode per cell by two-layer DNA seqFISH+ in NMuMG cells (right). *n* = 100,049 and 31,171 for on- and off-target barcodes, respectively. **e**, The distribution of mean on-target barcode counts per cell grouped by chromosome identities by two-layer DNA seqFISH+ in mESCs (top) and NMuMG cells (bottom). The differences in detection efficiency between autosomal and X chromosomal loci reflect the identities of male mESCs and female NMuMG cells. We note chromosome 19 in NMuMG cells could be trisomy because of the 43.0% greater average barcode counts per cell than those in the other chromosomes. *n* = 100,049 loci in total. **f**, Heatmap comparing average spatial distances of pairs of loci between two-layer DNA seqFISH+ (upper right) and previous DNA seqFISH+^7^ (lower left) at the DNA seqFISH+ 1-Mb resolution loci in Chr4 in mESCs. *n* = 149 loci. **g**, Pearson correlation of mean spatial distances of pairs of intra-chromosomal loci between two-layer DNA seqFISH+ and previous DNA seqFISH+^7^ at the DNA seqFISH+ 1-Mb resolution 25-kb loci across the genome in mESCs. **h**, Pearson correlation of mean spatial distances of pairs of intra-chromosomal loci between two biological replicates of two-layer DNA seqFISH+ using the same loci in **g** in mESCs. *n* = 159,397 pairs that were detected in both measurements in **g**, **h**. **i**, Representative visualization of eigenvectors between Hi-C^20, 67^ (top) and two-layer DNA seqFISH+ (bottom) for mESCs and NMuMG cells, confirming the highly concordant compartment organization between the measurements. **j**, Spearman correlation of eigenvectors between Hi-C and DNA seqFISH+ across the mouse genome for mESCs (top) and NMuMG cells (bottom). *n* = 23,547, 23,582 loci that were commonly profiled between the measurements for mESCs and NMuMG cells. 100 kb binning was used in **i**, **j**. **k**, Heatmap comparing median spatial distances of pairs of loci between two-layer DNA seqFISH+ (upper right) and previous DNA seqFISH+^7^ (lower left) at the previously profiled 25-kb resolution loci in Chr3, Chr13, and Chr14 in mESCs. *n* = 60 loci for each chromosome. Fluorescent channels used for two-layer DNA seqFISH+ loci are shown (top). We note that boundaries between loci in different fluorescent channels may be due to the differences in the efficiency of co-detecting pairs of loci across blocks (Method). **l**, Pearson correlation of median spatial distances of pairs of intra-chromosomal loci between two-layer DNA seqFISH+ and previous DNA seqFISH+^7^ at previously profiled 25-kb resolution loci at the selected regions in Chr1-19, X in mESCs. **m**, Pearson correlation of median spatial distances of pairs of intra-chromosomal loci between two biological replicates of two-layer DNA seqFISH+ using the same 25-kb loci in **l** in mESCs. *n* = 35,400 pairs that were commonly profiled between the measurements in **l**, **m**. *n* = 1,076 cells from two biological replicates of mESCs and *n* = 384 cells from one biological replicate of NMuMG cells in **b**-**m**.

**Extended Data Fig. 4.**
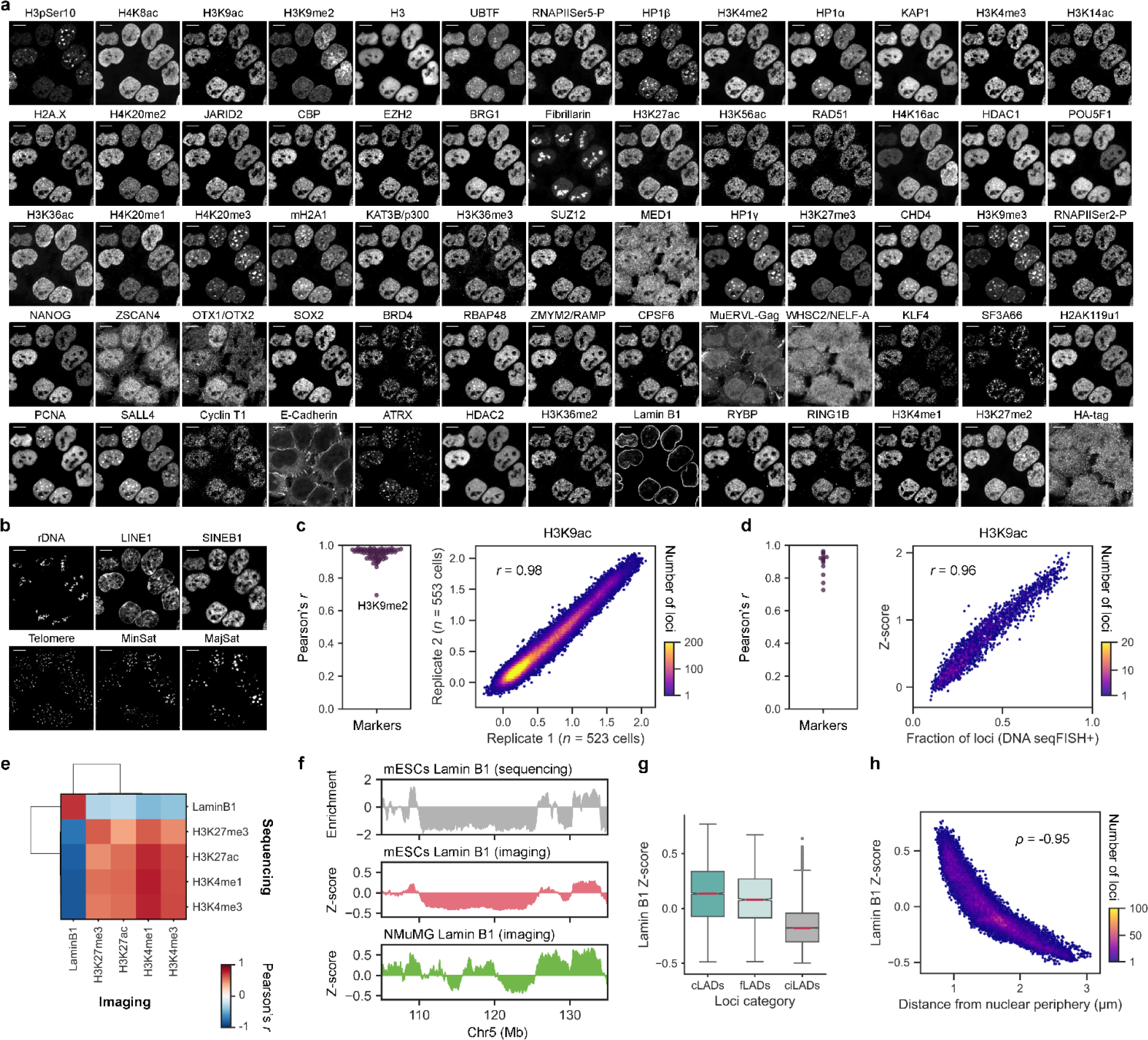
**Validation of imaging-based chromatin profiling across the mouse genome. a, b**, Representative images by sequential immunofluorescence (*n* = 65 markers) in **a** and DNA FISH of repetitive elements (*n* = 6 markers) in **b** in mESCs. **c**, Pearson correlation of imaging-based chromatin profiling for each marker (*n* = 69 markers in **a**, **b**, except E-Cadherin and HA-tag that do not stain nucleus) across the genome between two biological replicates in mESCs (left), showing H3K9me2 as an outlier, and corresponding density plot for individual loci (*n* = 100,049 loci) for H3K9ac as an example (right). **d**, Pearson correlation of imaging-based chromatin profiling for each marker (*n* = 13 markers) between two-layer DNA seqFISH+ and previous DNA seqFISH+^7^ at the DNA seqFISH+ 1-Mb resolution 25-kb loci across the genome in mESCs (left) and corresponding density plot for individual loci (*n* = 2,460 loci) for H3K9ac as an example (right). **e**, Pearson correlation of imaging-based chromatin profiling and sequencing-based chromatin profiling (DamID for Lamin B1^69^, CUT&Tag for other markers^70^) across the genome in mESCs. We note that imaging-based chromatin profiling measures spatial proximity between DNA loci and subnuclear structures, while sequencing-based profiling captures molecular interactions between genomic DNA and antibodies of interest. *n* = 25,110 loci that were commonly profiled between the measurements. **f**, Representative visualization of Lamin B1 chromatin profiles by DamID^69^ and two-layer DNA seqFISH+ in mESCs and NMuMG cells (from top to bottom), representing similar patterns between mESC datasets. **g**, Comparison of Lamin B1 enrichment among loci with different categories as constitutive lamina- associated domains (cLADs), facultative LADs (fLADs), and constitutive inter-LADs (ciLADs)^71, 72^ (*n* = 8,582, 2,908, and 10,239 loci), confirming higher Lamin B1 enrichments at LADs (cLADs and fLADs) than those at ciLADs. In box plots, the center lines for the median, boxes for the interquartile range, whiskers for values within 1.5 times the interquartile range, and points for outliers. **h**, Pearson correlation of imaging-based chromatin profiling of Lamin B1 and spatial distance from nuclear periphery in mESCs. Lamin B1 enriched DNA loci are physically closer to the nuclear periphery as expected. *n* = 25,148 loci. 100 kb binning was used in **e**-**h**. *n* = 1,076 cells from two biological replicates of mESCs and *n*= 384 cells from one biological replicate of NMuMG cells.

**Extended Data Fig. 5.**
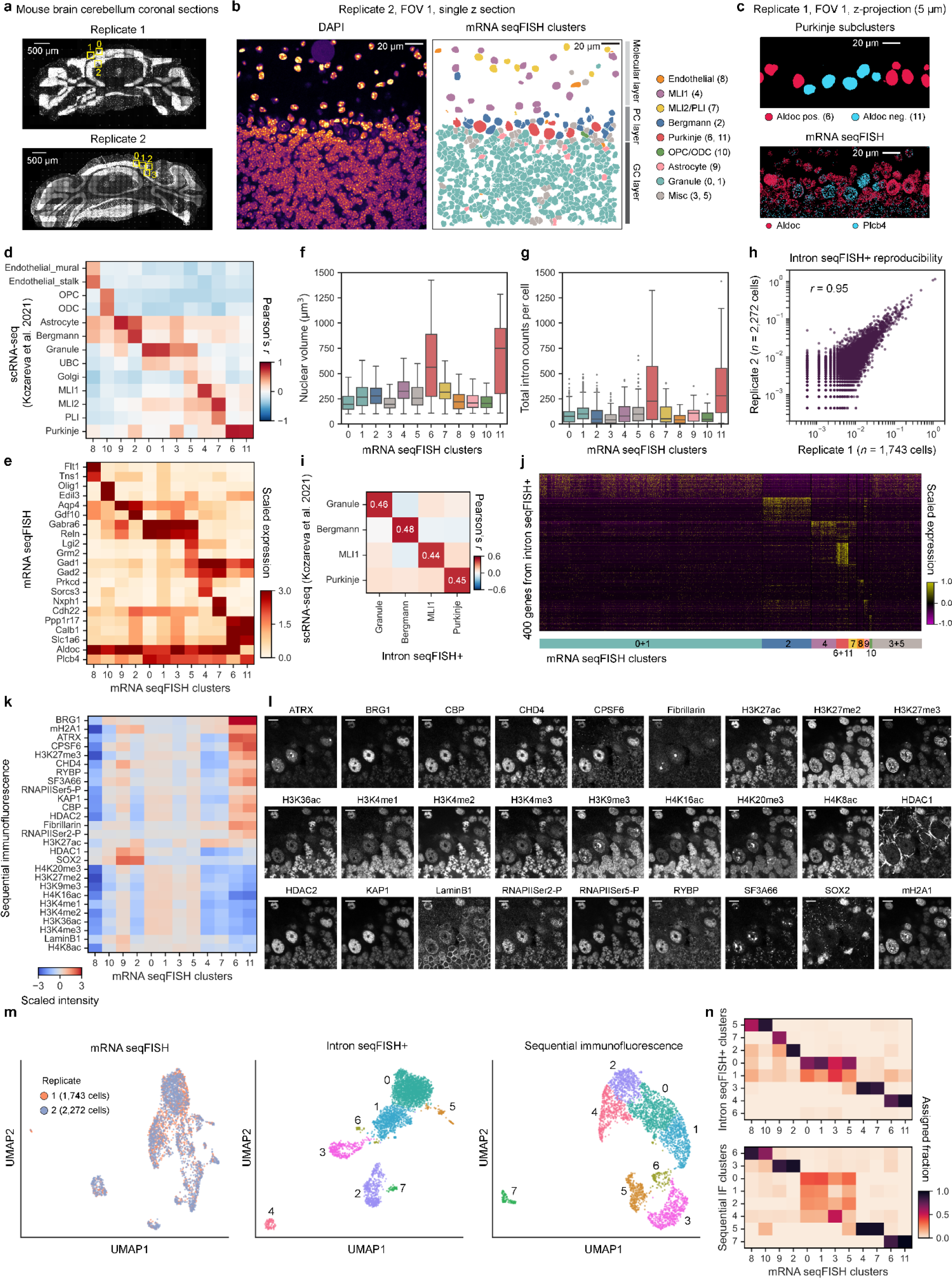
Validation and characterization of spatial multi-omics measurements in the **adult mouse brain cerebellum. a**, Large-field DAPI images of the adult mouse brain cerebellum coronal sections from two biological replicates. Yellow boxes represent unique fields of view (FOVs) imaged in each biological replicate. **b**, Representative images from a single z-section in one field of view for DAPI staining (left) and spatial distribution of cell clusters computed from mRNA seqFISH profiles (right). The cell type annotation for each cell cluster (0-11) was determined by the comparison with the single-nucleus RNA sequencing dataset^26^ shown in **d**. **c**, Representative images of spatial distribution of cell clusters for Purkinje cell subtypes (top) and corresponding marker gene expression in the nucleus (bottom), reflecting the known patterns of parasagittal stripes between Aldoc positive and negative Purkinje cells in the mouse cerebellum^26, 73, 74^. **d**, Comparison of cell clusters (0-11) defined by mRNA seqFISH and cell types identified by the single-nucleus RNA sequencing^26^. Based on the degree of Pearson correlation, we annotated our mRNA seqFISH clusters as shown in **b**. **e**, Marker gene expression profiles in each cell cluster. Those include previously characterized marker genes^26, 74^ such as Flt1 in Endothelial cells (cluster 8), Olig1 in oligodendrocyte precursor cells and oligodendrocytes (cluster 10), Aqp4 in Astrocytes (cluster 9), Gdf10 in Bergmann glia (cluster 2), Gabra6 in Granular cells (clusters 0 and 1), Sorcs3 in MLI1 (cluster 4), Nxph1 in MLI2/PLI (cluster 7), and Ppp1r17 in Purkinje cells (clusters 6 and 11, which can be further divided by Aldoc and Plcb4 as shown in **c**). **f**, Distribution of nuclear volume for cells in each cell cluster. **g**, Distribution of total intron counts per cell in each cell cluster by intron seqFISH+. In box plots, the center lines for the median, boxes for the interquartile range, whiskers for values within 1.5 times the interquartile range, and points for outliers in **f**, **g**. **h**, Reproducibility of intron seqFISH+ from two biological replicates with the adult mouse brain cerebellum. *n* = 17,849 genes that were detected at least once in each replicate. **i**, Comparison of gene expression profiles by intron seqFISH+ and single-nucleus RNA sequencing^26^ in four major cell types showed overall high consistency of cell-type specific gene expression programs. **j**, Heatmap of nascent gene expression profiles of differentially expressed genes by intron seqFISH+ across single cells grouped by cell clusters defined by mRNA seqFISH. **k**, Heatmap of sequential immunofluorescence intensity profiles across cell clusters defined by mRNA seqFISH. Note that HDAC1 and SOX2 were expressed in Bergmann glia (cluster 2) and astrocyte (cluster 9), consistent with previous studies^75, 76^. **l**, Representative sequential immunofluorescence images from a single z-section in the adult mouse brain cerebellum. **m**, UMAP-embedding of cells colored by two biological replicates (left) and each cell cluster defined by intron seqFISH+ profiles (middle) or sequential immunofluorescence intensity profiles (right). **n**, Comparison of cell clusters defined by mRNA seqFISH and intron seqFISH+ (top) or sequential immunofluorescence (bottom), representing robust identification of similar cell clusters regardless of the measurement modalities in the adult mouse brain cerebellum. *n* = 4,015 cells (*n* = 1,504, 832, 518, 357, 263, 164, 113, 88, 76, 56, 29, 15 cells from mRNA seqFISH cluster 0 to 11) from two biological replicates of the adult mouse brain cerebellum.

**Extended Data Fig. 6.**
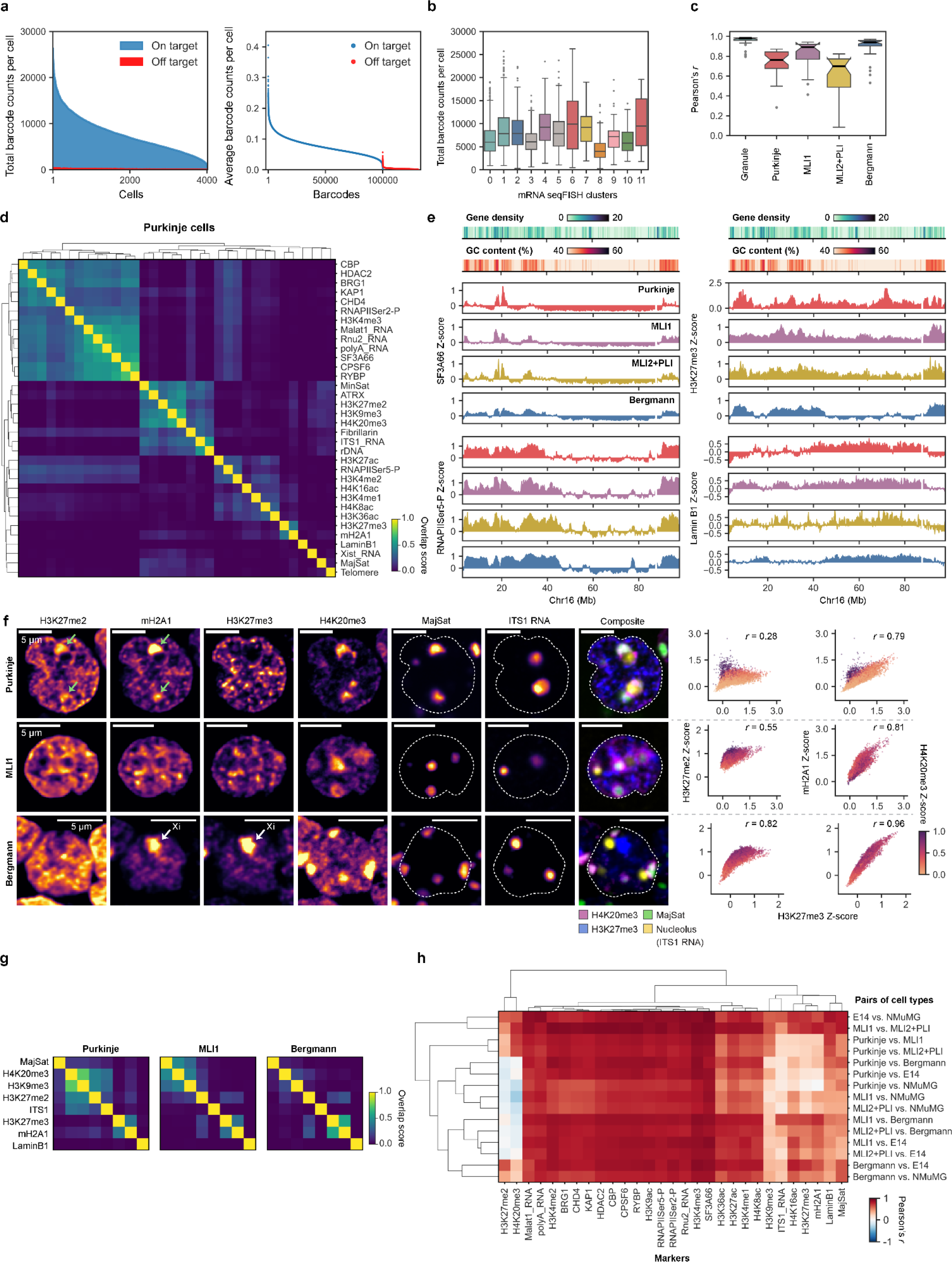
Additional validation and characterization of spatial multi-omics measurements in the adult mouse brain cerebellum. **a**, Total on- and off-target barcodes detected per cell by DNA seqFISH+ in the mouse cerebellum (left). Average counts per each on- and off-target barcode per cell by DNA seqFISH+ in the mouse cerebellum (right). *n* = 100,049 and 31,171 for on- and off-target barcodes, respectively. **b**, Total on-target barcodes detected per cell by DNA seqFISH+ in each transcriptionally-defined cell cluster in the mouse cerebellum. *n* = 1,504, 832, 518, 357, 263, 164, 113, 88, 76, 56, 29, 15 cells from mRNA seqFISH cluster 0 to 11. **c**, Pearson correlation of imaging-based chromatin profiling across the genome for each marker (*n* = 26 markers) in each cell type between two biological replicates of the adult mouse brain cerebellum. In box plots, the center lines for the median, boxes for the interquartile range, whiskers for values within 1.5 times the interquartile range, and points for outliers in **b**, **c**. **d**, Quantification of overlap of top 5% genomic loci associated with each marker between pairs of markers in Purkinje cells, largely separating active and repressive chromatin markers. **e**, Genomic features and representative imaging-based chromatin profiling with four markers in each cell type in chromosome 16. **f**, Representative raw immunofluorescence images for various repressive markers from a single z-section for each cell type (left). Purkinje-specific pericentromeric staining by H3K27me2 and mH2A1 is highlighted by green arrows. The intense mH2A1 and H3K27me3 clusters visible in Bergmann glia highlighted by white arrows represent the inactive X chromosome (Xi) territory^28^ in the female mouse cerebellum section. Scatter plots for individual loci show the relationship of each repressive marker enrichment in each cell type (right). **g**, Degree of overlap of top 5% enriched loci between pairs of repressive markers in each cell type. **h**, Degree of similarity of chromatin profiles between pairs of cell types by Pearson correlation, including cell lines (mESCs and NMuMG cells). 200 kb binning (*n* = 12,562 loci in total) was used for the analysis in **c**-**h**. *n* = 4,015 cells from two biological replicates of the adult mouse brain cerebellum.

**Extended Data Fig. 7.**
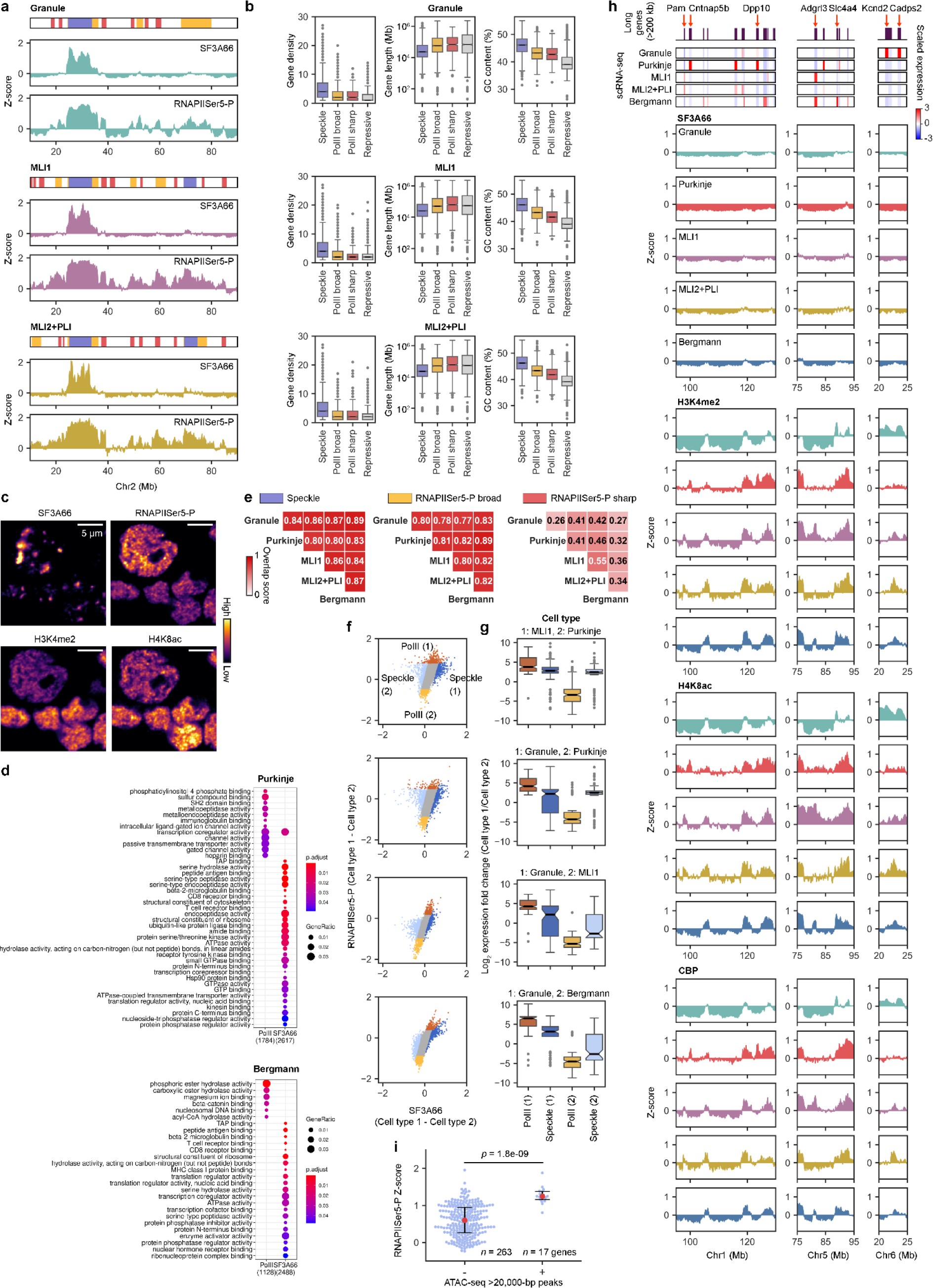
**Additional characterization of distinct active subnuclear compartments. a**, Representative classification of three types of active domains (nuclear speckle, RNAPIISer5-P broad and sharp) along with the chromatin profiling of SF3A66 (nuclear speckle marker) and RNAPIISer5-P across cell types. **b**, Comparison of genomic features across different active domains in each cell type. *n* = 2,531, 1,789, 458, 4,220 loci (Granule), *n* = 2,533, 2,060, 1,228, 4,654 loci (MLI1), and *n* = 2,376 2,015 1,367 4,210 loci (MLI2+PLI) from left to right category. **c**, Representative raw images of active markers from a single z-section of the adult mouse brain cerebellum, showing distinct compartmentalization between nuclear speckles (SF3A66) and other active markers (e.g. RNAPIISer5-P, H3K4me2, H4K8ac) regardless of the cell types. **d**, GO term comparison between SF3A66 and RNAPIISer5-P associated genomic loci in each cell type, representing largely distinct enrichment. Similar observation of distinct pathway enrichments between speckle-associating and speckle-non-associating p53 target genes was previously observed in human cell lines^32^. **e**, Overlap of each active domain across cell types, showing more cell-type specific organization of RNAPIISer5-P sharp domains. **f**, Comparison of differential association of genomic loci with SF3A66 and RNAPIISer5-P between pairs of cell types. **g**, Comparison of differential mRNA expression^26^ between pairs of cell types at differentially associated loci with either PolII (RNAPIISer5-P) or Speckle (SF3A66) in each cell type. *n* = 150, 114, 198, 126 loci (MLI1 vs. Purkinje), *n* = 203, 136, 139, 146 loci (Granule vs. Purkinje), *n* = 164, 85, 143, 79 loci (Granule vs. MLI1), and *n* = 147, 96, 117, 87 loci (Granule vs. Bergmann) from left to right category. In box plots, the center lines for the median, boxes for the interquartile range, whiskers for values within 1.5 times the interquartile range, and points for outliers in **b**, **g**. **h**, Representative genomic regions with long genes (>200 kb) (top), corresponding mRNA expression^26^ (middle), and chromatin profiles of different active markers (bottom) in each cell type. Nuclear speckles marked by SF3A66 were not enriched at the differentially expressed long genes highlighted with red arrows (top). **i**, Comparison of Purkinje RNAPIISer5-P enrichment between long genes with the absence or presence of highly open chromatin regions defined by ATAC-seq >20 kb peaks in Purkinje cells^77^. The red dots for the median and error bars for the interquartile range. *p* values by two-sided Wilcoxon’s signed rank-sum test. 200 kb binning (*n* = 12,562 loci in total) was used for the analysis in **a**, **b**, **e**-**h**. *n* = 2,336, 128, 263, 88, and 518 cells for Granule, Purkinje, MLI1, MLI2+PLI, and Bergmann glia cells from two biological replicates of the adult mouse cerebellum.

**Extended Data Fig. 8.**
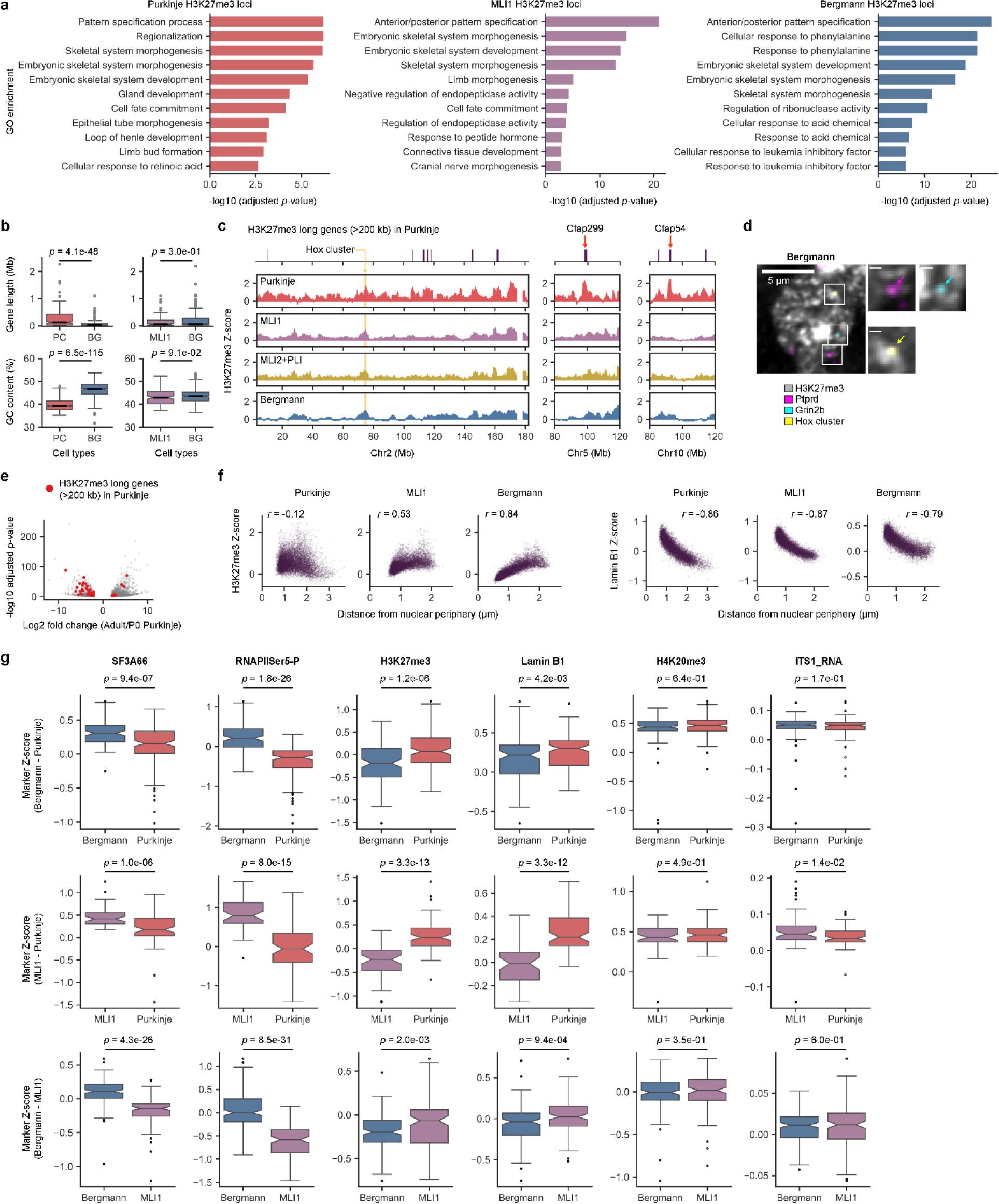
**Additional visualization and characterization of cell-type specific repressive subnuclear compartments. a**, The top eleven GO terms identified from H3K27me3 chromatin profiles in each cell type represent the enrichment of similar GO terms across cell types. **b**, Comparison of genomic features between loci that were differentially associated with H3K27me3 between two cell types. *n* = 595, 359 loci for Purkinje cells (PC) vs. Bergmann glia (BG) and *n* = 169, 491 loci for MLI1 vs. BG. **c**, Additional examples of H3K27me3 profiles across cell types along with Hox cluster in Chr2 and Purkinje-specific H3K27me3-associated long gene loci (top). **d**, Representative single cell visualization of genomic loci relative to H3K27me3 in Bergmann glia. Scale bars, 500 nm. **e**, Violin plot of expression changes between P0 and adult Purkinje cells^46^, representing the down-regulaiton of 36 out of 46 differentially expressed genes categorized as Purkinje-specific H3K27me3-associated long genes from the P0 to adult. **f**, Comparison of spatial distance from nuclear periphery and H3K27me3 or Lamin B1 chromatin profiles in each cell type. **g**, Comparison of differential association with each subnuclear marker at differentially expressed loci defined by intron seqFISH+ between pairs of major cell types. The increased association with SF3A66 at transcriptionally up-regulated loci was similarly observed in the adult mouse cerebral cortex^8^. *n* = 114, 129 loci for Bergmann glia vs. Purkinje cells, *n* = 48, 81 loci for MLI1 vs. Purkinje cells, and *n* = 151, 114 loci for Bergmann glia vs. MLI1. *p* values by two-sided Wilcoxon’s signed rank-sum test in **b**, **g**. In box plots, the center lines for the median, boxes for the interquartile range, whiskers for values within 1.5 times the interquartile range, and points for outliers in **b**, **g**. 200 kb binning (*n* = 12,562 loci in total) was used for the analysis in **a**-**c**, **f**, **g**. *n* = 128, 263, 88, and 518 cells for Purkinje, MLI1, MLI2+PLI, and Bergmann glia cells from two biological replicates of the adult mouse cerebellum.

**Extended Data Fig. 9.**
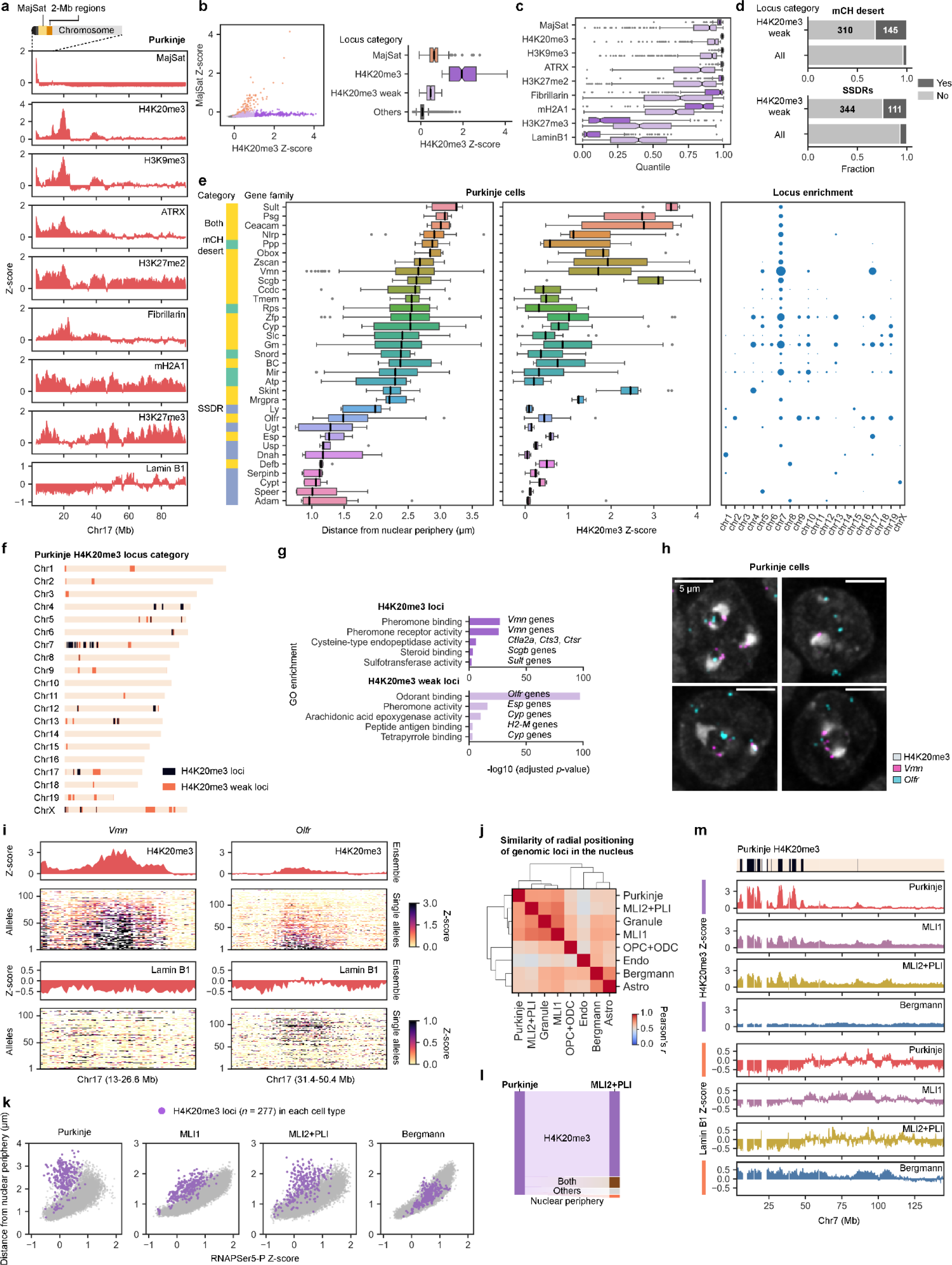
Additional characterization of H4K20me3 subnuclear compartments. a, Cartoon illustration of a mouse chromosome (top) and representative imaging-based chromatin profiling of repressive markers in Purkinje cells in chromosome 17 (bottom). **b**, Scatter plots of pairs of chromatin profiles (left), colored by the locus category of MajSat (orange), H4K20me3 (purple), H4K20me3 weak (light purple), or others (gray) and corresponding box plots showing H4K20me3 enrichment for loci in each category (right). **c**, Comparison of quantile of H4K20me3 and H4K20me3-weak category loci in **b** for each repressive marker, representing stronger enrichment of repressive markers around pericentromeric heterochromatin (MajSat, H4K20me3, H3K9me3, ATRX, H3K27me2, Fibrillarin, and mH2A1) at H4K20me3 loci (purple) over H4K20me3-weak loci (light purple). In contrast, two repressive markers (H3K27me3 and Lamin B1) are more depleted at H4K20me3 loci compared to H4K20me3-weak loci. **d**, Barplots comparing the locus characteristics such as mCH desert^48^ and SSDRs^50, 51^ between H4K20me3-weak loci (*n* = 455) and all 200-kb loci (*n* = 12,562) in Purkinje cells. **e**, Gene family characteristics either mCH desert^48^, SSDRs^50, 51^, or both (left) and their radial positioning relative to nuclear periphery and H4K20me3 enrichment (middle), as well as their enrichments across chromosomes (right). Only a subset of genomic loci annotated with those categories with corresponding gene family names were included in this analysis. In box plots, the center lines for the median, boxes for the interquartile range, whiskers for values within 1.5 times the interquartile range, and points for outliers in **b**, **c**, **e**. **f**, H4K20me3 and H4K20me3-weak loci (*n* = 236, 455, respectively) distribution across chromosomes in Purkinje cells. **g**, GO term comparison between H4K20me3 and H4K20me3-weak associated genomic loci in Purkinje cells revealed enrichment of distinct gene families at each category. **h**, Additional visualization of H4K20me3-enriched *Vmn* and *Olfr* gene family loci overlaid with H4K20me3 staining with a maximum z-projection of four sections in Purkinje cells. **i**, Additional examples of ensemble-averaged and single allele chromatin profiles sorted by H4K20me3 enrichment from bottom to top in Purkinje cells. **j**, Comparison of ensemble-averaged radial positioning of genome-wide DNA loci in the nucleus across cell types separates neurons and glial cells. **k**, Comparison between RNAPIISer5-P enrichment and spatial distance from nuclear periphery for individual loci. Top 5% H4K20me3-enriched loci (purple, *n* = 277) in each cell type tend to localize in the nuclear interiors yet excluded from RNAPIISer5-P in neurons (Purkinje, MLI1, and MLI2/PLI) but not in Bergmann glia cells. Those neuron-specific outlier loci found here were not observed in human cell culture^30^. **l**, Transition of H4K20me3-enriched loci from Purkinje to MLI2/PLI cells showed largely conserved H4K20me3-associated loci between neuronal cell types. *n* = 252 loci. **m**, The H4K20me3-enriched regions in Purkinje cells (top) and H4K20me3 and Lamin B1 chromatin profiles across cell types in Chr7 (bottom). 200 kb binning (*n* = 12,562 loci in total) was used for the analysis in **a**-**g**, **i**-**m**. *n* = 4,015 cells from two biological replicates of the adult mouse brain cerebellum.

**Extended Data Fig. 10.**
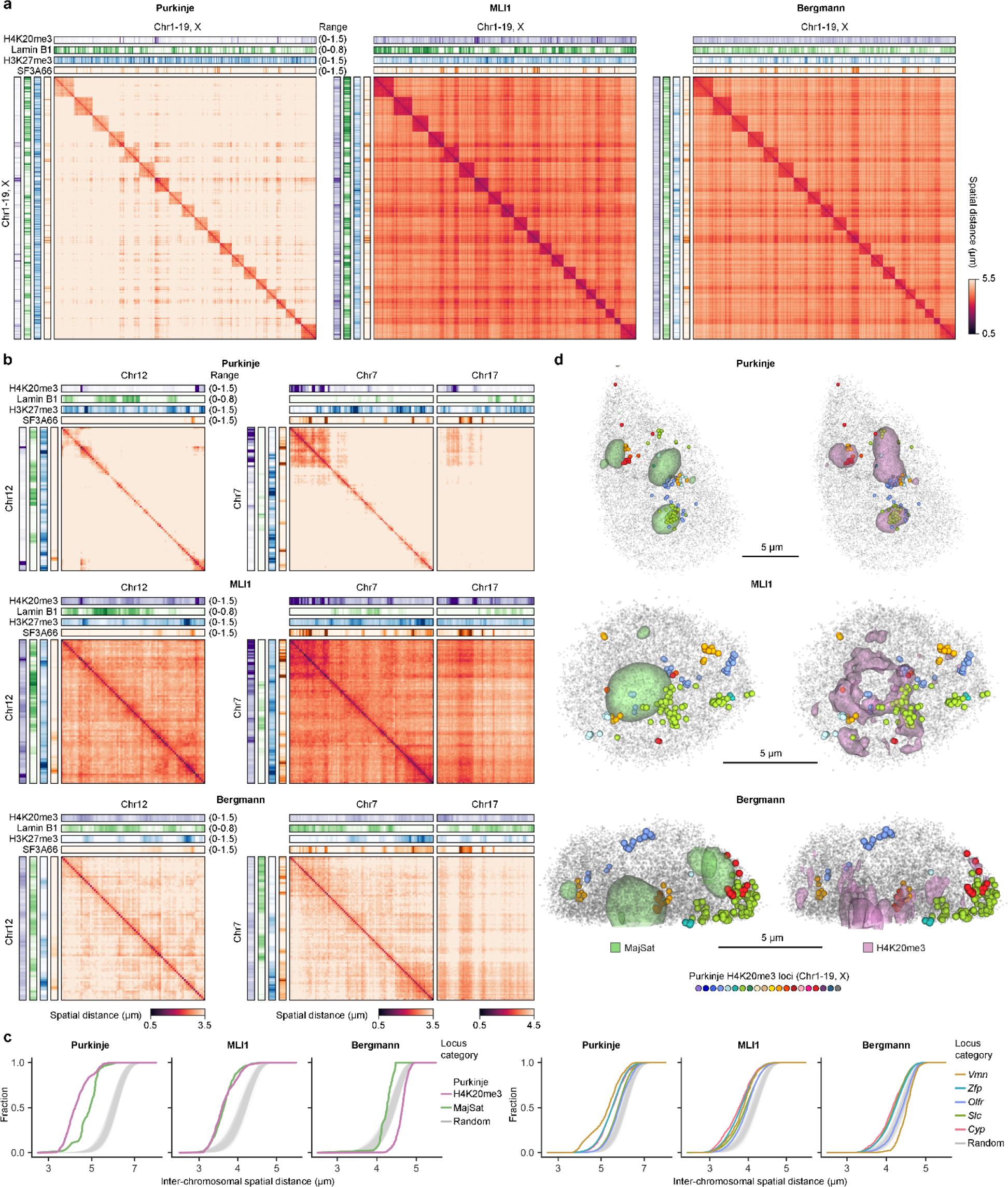
**Additional visualization and characterization of 3D genome organization with subnuclear structures. a**, Ensemble-averaged spatial distances between pairs of genomic loci along with chromatin profiling from each cell type across chromosomes (Chr1-19, X). **b**, Additional examples of ensemble-averaged spatial distances between pairs of genomic loci along with chromatin profiling from each cell type at specific chromosomes. **c**, Cumulative distribution of inter-chromosomal distances between pairs of loci with top 5% association to MajSat or Purkinje-specific H4K20me3 characterized in Fig. 5c (left) or from loci associated with specific gene families (right) compared to random pairs of loci (*n* = 1,000 trials). **d**, Representative 3D images of MajSat or H4K20me3 staining and chromosomal loci in each cell type. Identified Purkinje H4K20me3 loci highlighted with colors tend to colocalize at H4K20me3 territories in Purkinje and MLI1 cells, but localize at the nuclear periphery in Bergmann glia cells. 1.5 Mb binning (*n* = 1,678 loci in total), grouped by the chromosome paint block barcodes, was used for the analysis in **a**-**c**. *n* = 128, 263, and 518 cells for Purkinje, MLI1, and Bergmann glia cells from two biological replicates of the adult mouse cerebellum.

## Supplementary Tables

**Supplementary Table 1**: A list of genomic coordinates for the 100,049 DNA loci used in two-layer DNA seqFISH+ with barcoding information

**Supplementary Table 2**: A list of primary antibodies used in this study.

**Supplementary Table 3**: A summary of 4,015 cells profiled in the adult mouse cerebellum in this study with information of replicate ID, fields of view (FOV) ID for each replicate, cell ID for each FOV and each replicate, mRNA seqFISH cluster (0-11), centroid voxel coordinates of the nucleus, and nuclear volume (μm^3^).

